# Cross-family interactions of vascular endothelial growth factors and platelet-derived growth factors on the endothelial cell surface: A computational model

**DOI:** 10.1101/2025.02.27.640640

**Authors:** Yunjeong Lee, Yingye Fang, Shobhan Kuila, Princess I. Imoukhuede

**Author notes:** Corresponding author (PII).

## Abstract

Angiogenesis, the formation of new vessels from existing vessels, is mediated by vascular endothelial growth factor (VEGF) and platelet-derived growth factor (PDGF). Despite discoveries supporting the cross-family interactions between VEGF and PDGF families, sharing the binding partners between them makes it challenging to identify growth factors that predominantly affect angiogenesis. Systems biology offers promises to untangle this complexity. Thus, in this study, we developed a mass-action kinetics-based computational model for cross-family interactions between VEGFs (VEGF-A, VEGF-B, and PlGF) and PDGFs (PDGF-AA, PDGF-AB, and PDGF-BB) with their receptors (VEGFR1, VEGFR2, NRP1, PDGFRα, and PDGFRβ). The model, parametrized with our literature mining and surface resonance plasmon assays, was validated by comparing the concentration of VEGFR1 complexes with a previously constructed angiogenesis model. The model predictions include five outcomes: 1) the percentage of free or bound ligands and 2) receptors, 3) the concentration of free ligands, 4) the percentage of ligands occupying each receptor, and 5) the concentration of ligands that is bound to each receptor. We found that at equimolar ligand concentrations (1 nM), PlGF and VEGF-A were the main binding partners of VEGFR1 and VEGFR2, respectively. Varying the density of receptors resulted in the following five outcomes: 1) Increasing VEGFR1 density depletes the free PlGF concentration, 2) increasing VEGFR2 density decreases PDGF:PDGFRα complexes, 3) increased NRP1 density generates a biphasic concentration of the free PlGF, 4) increased PDGFRα density increases PDGFs:PDGFRα binding, and 5) increasing PDGFRβ density increases VEGF-A:PDGFRβ. Our model offers a reproducible, fundamental framework for exploring cross-family interactions that can be extended to the tissue level or intracellular molecular level. Also, our model may help develop therapeutic strategies in pathological angiogenesis by identifying the dominant complex in the cell signaling.

**Author summary:** New blood vessel formation from existing ones is essential for growth, healing, and reproduction. However, when this process is disrupted—either too much or too little—it can contribute to diseases such as cancer and peripheral arterial disease. Two key families of proteins, vascular endothelial growth factors (VEGFs) and platelet-derived growth factors (PDGFs), regulate this process. Traditionally, scientists believed that VEGFs only bind to VEGF receptors and PDGFs to PDGF receptors. However, recent findings show that these proteins can interact with each other’s receptors, making it more challenging to understand and control blood vessel formation. To clarify these complex interactions, we combined computer modeling with biological data to map out which proteins bind to which receptors and to what extent. Our findings show that when VEGFs and PDGFs are present in equal amounts, VEGFs are the primary binding partners for VEGF receptors. We also explored how changes in receptor levels affect these interactions in disease-like conditions. This work provides a foundational computational model for studying cross-family interactions, which can be expanded to investigate tissue-level effects and processes inside cells. Ultimately, our model may help develop better treatments for diseases linked to abnormal blood vessel growth by identifying key protein-receptor interactions.

## Introduction

Angiogenesis, the process of new microvessel formation from existing vessels, is involved in physiological and pathological processes. Vascular endothelial growth factors (VEGFs), the main regulators of angiogenesis, promote the proliferation, migration, and vascular permeability of endothelial cells [1,2]. Another important supporter of vascular development is the platelet-derived growth factors (PDGFs), which enable vascular maturation via pericyte recruitment [3].

Five VEGF isoforms have been identified as important in human vascular development: VEGF-A, VEGF-B, VEGF-C, VEGF-D, and placental growth factor (PlGF). The VEGF receptor family includes three receptors— VEGFR1, VEGFR2, and VEGFR3—which can exist in two forms: transmembrane receptors, located either on the plasma membrane or within intracellular organelles [4], and soluble receptors, which are secreted but may also localize intracellularly depending on the splice variant [5–8]. Neuropilin-1 (NRP1) serves as a co-receptor that stabilizes ligand binding to VEGFRs and potentiate the signaling by forming VEGF:VEGFR:NRP1 complexes [9]. These numerous possibilities for VEGF interacting with VEGFRs and co-receptors lead to the activation of multiple downstream signaling pathways that induce the hallmark angiogenic responses: endothelial cell migration, proliferation, survival, and ultimately anastomoses [2,10,11].

The selective binding results in differential outcomes (Fig 1A). VEGF-A165a is considered the most potent VEGF family member in angiogenesis (herein described as VEGF-A). Alternative splicing generates several proangiogenic VEGF-A isoforms, including VEGF-A121, VEGF-A165a, VEGF-A189, and VEGF-A206, and several modulatory isoforms such as VEGF-A165b [12,13]. While VEGF-A can bind to both VEGFR1 and VEGFR2, the complex of VEGF-A and VEGFR2 is the most well-known key factor inducing angiogenesis. VEGFR1 is often described as a decoy receptor because the tyrosine kinase activity and phosphorylation level of VEGFR1 induced by VEGF is weaker than that of VEGFR2, despite having a stronger affinity for VEGF than to VEGFR2 [14,15]. This decoy nomenclature is also attributed to VEGFR1 knockout mouse studies, which demonstrated disorganized vasculature possibly due to excessive angiogenesis induced by the lack of VEGF-A:VEGFR1 signaling [14,16]. VEGF-B family includes two isoforms: VEGF-B167 and VEGF-B186, but VEGF-B167 (hereafter VEGF-B) is 5–10 times more expressed than VEGF-B186 in normal tissues [17]. Both VEGF-B isoforms can bind to VEGFR1 and NRP1 although VEGF-B186 should be proteolytically processed to bind NRP1 [18]. The VEGF-B family was first established to regulate vessel survival rather than vessel growth [19,20]. Recent studies have revealed that their complexes with VEGFR1 and/or NRP1 generate signals to regulate fatty acid transport into the myocardium, skeletal muscles, and brown adipose tissue [21]. VEGF-C and VEGF-D are involved in both angiogenesis and lymphangiogenesis [22], but they mainly regulate lymphangiogenesis via activating VEGFR3 [23]. PlGF is mainly expressed in the placenta and to a lesser extent in other tissues in human bodies [24]. Human tissues express four isoforms: PlGF-1, PlGF-2, PlGF-3, and PlGF-4 [24,25], while mouse tissue expresses only one isoform, PlGF-2 [26]. PlGF augments VEGF-A-induced angiogenic signaling by competitively binding to VEGFR1 [27]. Also, PlGF:VEGFR1 complex induces proangiogenic signaling by itself or by intermolecular crosstalk between VEGFR1 and VEGFR2 [28]. Although the PlGF is a VEGF family member, it is called “placental” growth factor since it was initially cloned from a human placental cDNA library [29,30].

**Fig 1.**
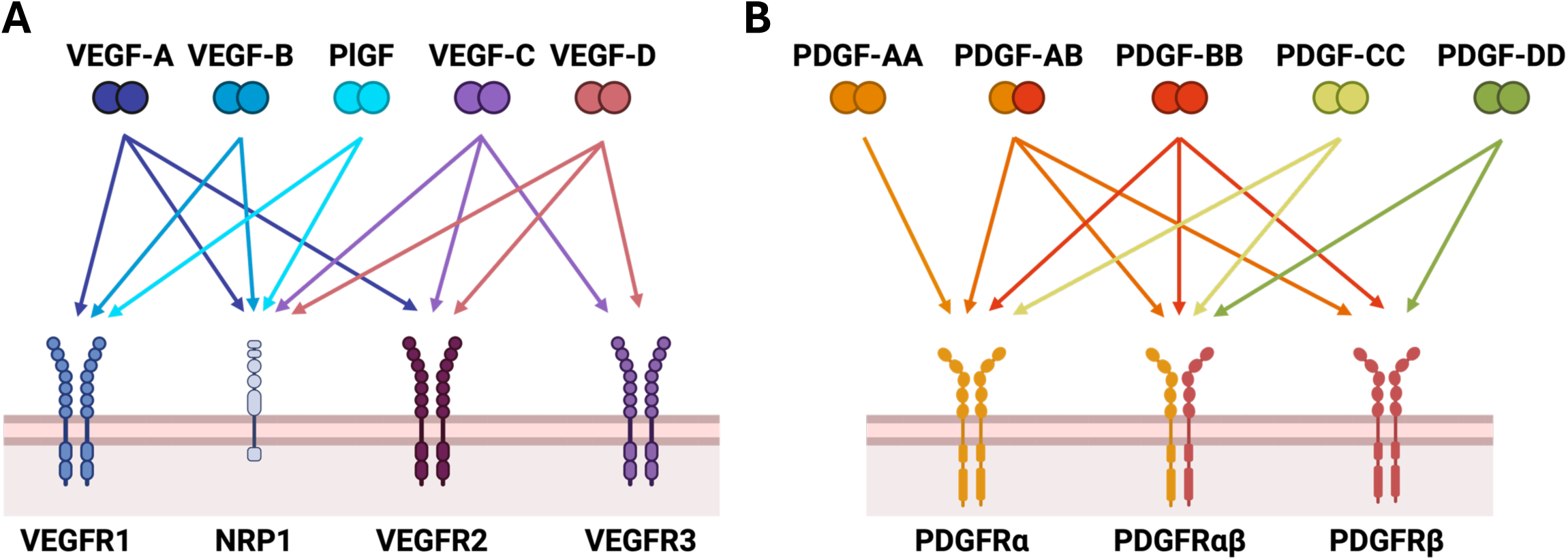
VEGF and PDGF binding to their receptors. (A) Five VEF isoforms bind to VEGFR families and NRP1. VEGF-A binds to VEGFR1, VEGFR2, and NRP1, while VEGF-B and PlGF binds to VEGFR1 and NRP1. VEGF-C and VEGF-D binds to VEGFR2, VEGFR3, and NRP1. (B) Five PDGF isoforms interact with three PDGFRs: PDGFRα, PDGFRαβ, and PDGFRβ. PDGFRα binds to all PDGFs except for PDGF-DD, while PDGFRαβ binds to all PDGFs except for PDGF-AA. PDGFRβ binds to only PDGF-BB and PDGF-DD.

The PDGF family includes four homodimeric proteins: PDGF-AA, PDGF-BB, PDGF-CC, and PDGF-DD, in addition to one heterodimeric protein, PDGF-AB (Fig 1B). PDGF family members selectively bind to two homodimerized receptors, PDGFRα and PDGFRβ, and one heterodimerized receptor, PDGFRαβ. While PDGFRα can bind to all PDGF members except PDGF-DD, PDGFRβ can bind to PDGF-AB, PDGF-BB, and PDGF-DD only, and PDGFRαβ binds to all PDGFs except PDGF-AA. Although all PDGF is involved in angiogenesis [31], among PDGF isoforms consisting of two subunits A or B, PDGF-AA has been demonstrated to exhibit a lower potency in promoting angiogenesis than PDGF-AB and PDGF-BB [32]. PDGF-BB has been shown to enhance the migration of endothelial cells and facilitate the recruitment of pericytes and smooth muscle cells during angiogenesis [33–35]. Additionally, PDGF-CC promotes the proliferation and survival of endothelial cells, pericytes, and fibroblasts through PDGFRα activation [36]. PDGF-DD enhances angiogenic sprouting and increases the migration of pericytes by possibly mediating intercellular signaling between endothelial cells and pericytes [37]. However, PDGF-CC and PDGF-DD are secreted in inactive forms; thus, proteolytic cleavage is required before receptor binding [38–40].

The conventional perspective on VEGF and PDGF signaling has been a uni-family interaction: VEGF binds to VEGFRs only, and PDGF binds to PDGFRs. However, increasing evidence indicates cross-family interactions, i.e., VEGF and PDGF can bind to each other’s receptors: 1) VEGF-A binds to PDGFRs with binding affinity ranging from 0.3–26 nM while PDGFs bind to VEGFRs with 70 pM–0.53 µM binding affinities [41], 2) VEGF stimulates the migration of mesenchymal stem cells and fibroblasts by activating PDGFRα and PDGFRβ [42], and 3) VEGF-A dose-dependently activates PDGFRs [42] and can inhibit PDGF-induced PDGFR activation through competitive binding [43]. This evidence also supports the necessity of exploring cross-family interactions. Indeed, the administration of two growth factors, such as fibroblast growth factor-2 and PDGF-AB, has shown significantly enhanced angiogenesis in mouse corneal assay and rat ischemic hind-limb model compared to the administration of a single growth factor [32].

The complexity of competitive interactions between growth factors makes it challenging to identify growth factors that predominantly affect angiogenesis. Systems biology, incorporating computational models and experimental data, is well-suited to unmixing the complexity inherent to multi-growth factor-receptor interactions. The most well-established computational models for angiogenesis are found within the field of tumor research. Several tumor models have: 1) investigated the distribution of VEGF-A isoforms, i.e., the proportion of VEGF-A that are free, bound to receptors, or bound to extracellular matrix, in tumors and 2) evaluated the effect of treatment targeting VEGF-A on tumor angiogenesis [44,45]. However, these models assumed uni-family interactions, not cross-family interactions. Mamer et al. (2017) established the first computational model for cross-family interaction between PDGFs and VEGFR2 while also considering VEGF-A:VEGFR2 binding [41]. Our model builds upon this foundational cross-family model by considering VEGF-A:PDGFR binding. Thus, our goal here is two-fold: 1) establish a computational model for cross-family interactions between VEGF and PDGF families on the endothelial cell surface and 2) understand the significance of cross-family interactions and its extent by identifying the steady-state distribution of VEGF and PDGF isoforms occupying each receptor. To achieve this goal, we established a system of coupled ordinary differential equations by assuming the mass-action kinetics. We validated the model by comparing to previously verified computational model for VEGF and PlGF treatment on *in vitro* endothelial cells. We varied receptor density on an endothelial cell and investigated their effects on the ligand distribution. Our computational model offers foundational insight into the significance of cross-family interactions while also enabling future control of this newly recognized interaction and signaling mechanism.

## Method

### Computational model

We developed a computational model to investigate cross-family interactions on the endothelial cell membrane. This model focuses on the ligand-receptor interactions between VEGFs, PDGFs, and their receptors. Six ligands were included: VEGF-A, VEGF-B, PlGF-2 (PlGF), PDGF-AA, PDGF-AB, and PDGF-BB. Only PlGF-2 was included since we established the model for the endothelial cell in mice adipose tissue and PlGF-2 is the only isoform that is expressed in mice [26]. Although VEGF-C and VEGF-D have some proangiogenic function [22], they are not established as the major signaling molecules in this process, and their primary roles are in lymphangiogenesis [23], as such, we did not include them in this model. Additionally, we did not model the cleavage process of growth factors, leading to the exclusion of PDGF-CC and PDGF-DD, which require proteolytic cleavage. The model incorporates five receptors: VEGFR1, VEGFR2, NRP1, PDGFRα, and PDGFRβ. Heterodimers were not considered due to lack of data on how heterodimers are part of cross-family interactions.

The binding properties of VEGF and PDGF isoforms with their receptors vary (Fig 2A): VEGF-A binds to all receptors, while VEGF-B and PlGF bind only to VEGFR1 and NRP1. All PDGF isoforms bind to VEGFR2; however, only PDGF-AA binds specifically to PDGFRα, while the other isoforms can bind to both PDGFRs. Furthermore, receptors can form complexes with or without ligands (Fig 2B): 1) VEGFR1 binds to NRP1 to generate a VEGFR1:NRP1 complex, which does not bind to growth factors; 2) VEGFR2, when bound to VEGF-A, also binds to NRP1, and vice versa. Overall, the model comprises 30 species, 40 reactions, and 30 ordinary differential equations (for detailed equations, please refer to Supplementary File).

**Fig 2.**
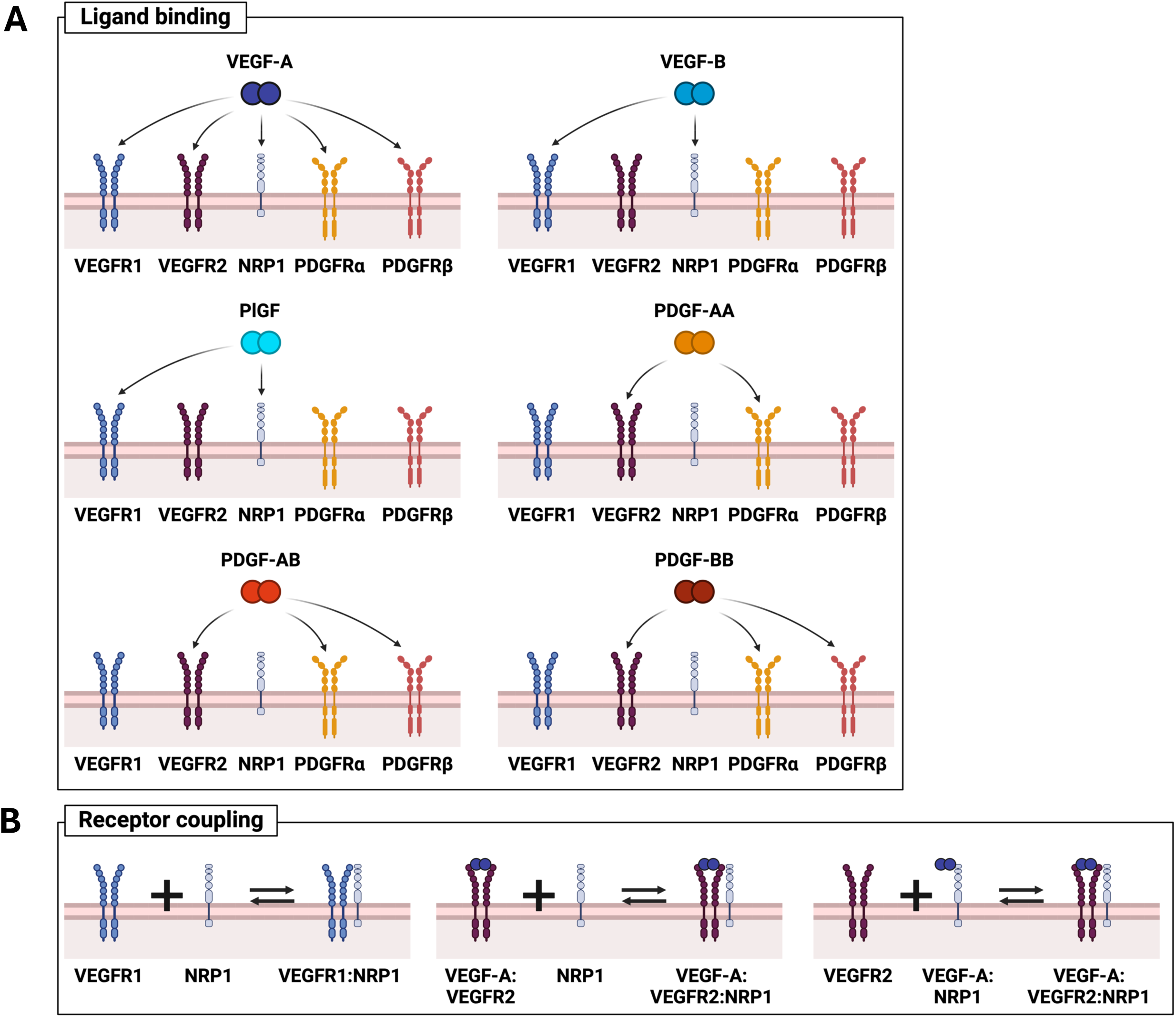
The schematic of cross-family signaling on the surface of an endothelial cell. (A) VEGF-A binds to all types of receptors: VEGFR1, VEGFR2, NRP1, PDGFR⍺, and PDGFRβ. VEGF-B and PlGF bind to VEGFR1 and NRP1. PDGF-AA, PDGF-AB, and PDGF-BB bind to VEGFR2 and PDGFR⍺ while PDGF-AB and –BB bind to PDGFRβ unlike PDGF-AA. (B) Receptors also form complex with or without ligand binding. VEGFR1 can form a complex with NRP1. VEGF-A:VEGFR2 complex can bind to NRP1, leading to a VEGF-A:VEGFR2:NRP1 complex. VEGF-A:NRP1 complex can bind to VEGFR2, forming a VEGF-A:VEGFR2:NRP1 complex, again.

### Model assumptions

The model assumed: 1) ligands are neither secreted nor degraded and 2) receptors are not internalized, recycled, degraded, or synthesized. The initial concentrations of ligands were assumed to be 1 nM (Table 1). This concentration was chosen because 1 nM of growth factors is commonly used for the stimulation of endothelial cells [46–48] and the 50 ng/ml of VEGF-A (around 1 nM for VEGF-A considering its molecular weight 45 kDa) has been reported to be optimal to endothelial cell signaling [49,50]. Although, it should be noted that serum concentrations of these growth factors differ (VEGF-A 2 pM, and PDGFs 9–350 pM [41]).

**Table 1.** Ligand concentration and receptor densities.

| Parameter | Ligand | Measured value | Model value | Reference |
| --- | --- | --- | --- | --- |
| Ligand concentration | VEGF-A | - | 1 nM | Assumed |
|  | VEGF-B | - | 1 nM | Assumed |
|  | PlGF | - | 1 nM | Assumed |
|  | PDGF-AA | - | 1 nM | Assumed |
|  | PDGF-AB | - | 1 nM | Assumed |
|  | PDGF-BB | - | 1 nM | Assumed |
| Receptor density | VEGFR1 | 1,600 rec/cell | 2.66 nM | Mouse Adipose ECs [51] |
|  | VEGFR2 | 4,100 rec/cell | 6.81 nM | Mouse Adipose ECs [51] |
|  | NRP1 | 44,100 rec/cell | 73.2 nM | HUVECs [52] |
| | PDGFRα | 5,000 rec/cell | 8.30 nM | Mouse Adipose ECs [51] |
| | PDGFRβ | 5,300 rec/cell | 8.80 nM | Mouse Adipose ECs [51] |

“rec/cell” represents the number of receptors on the endothelial cell surface. The unit, “rec/cell”, was converted by multiplying $1.66 \times 10^{-12}$ M/(rec/cell) ( $= 1/[\text{Avogadro number}] \times 1/[\text{endothelial cell volume}]$ ).
Abbreviation: endothelial cell (EC); human umbilical vein endothelial cells (HUVEC)

### Model parameterization

The parameters used in the model are presented in Tables 1 and 2. The densities of VEGFRs and PDGFRs on the endothelial cell surface were quantified from endothelial cells in adipose tissue [51]. NRP1 density measured in human umbilical vein endothelial cells with no ligand treatment was used [52]. The unit of receptor densities on the cell surface (rec/cell) was converted into molarity (M) using Avogadro’s number (6.02214×10^23^ molecule/mol) and the volume of an endothelial cell (1 pL) [53].

**Table 2.**
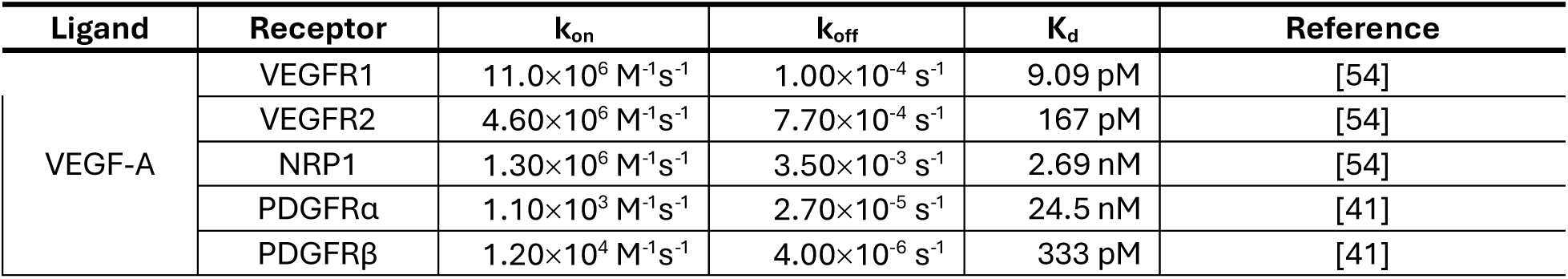
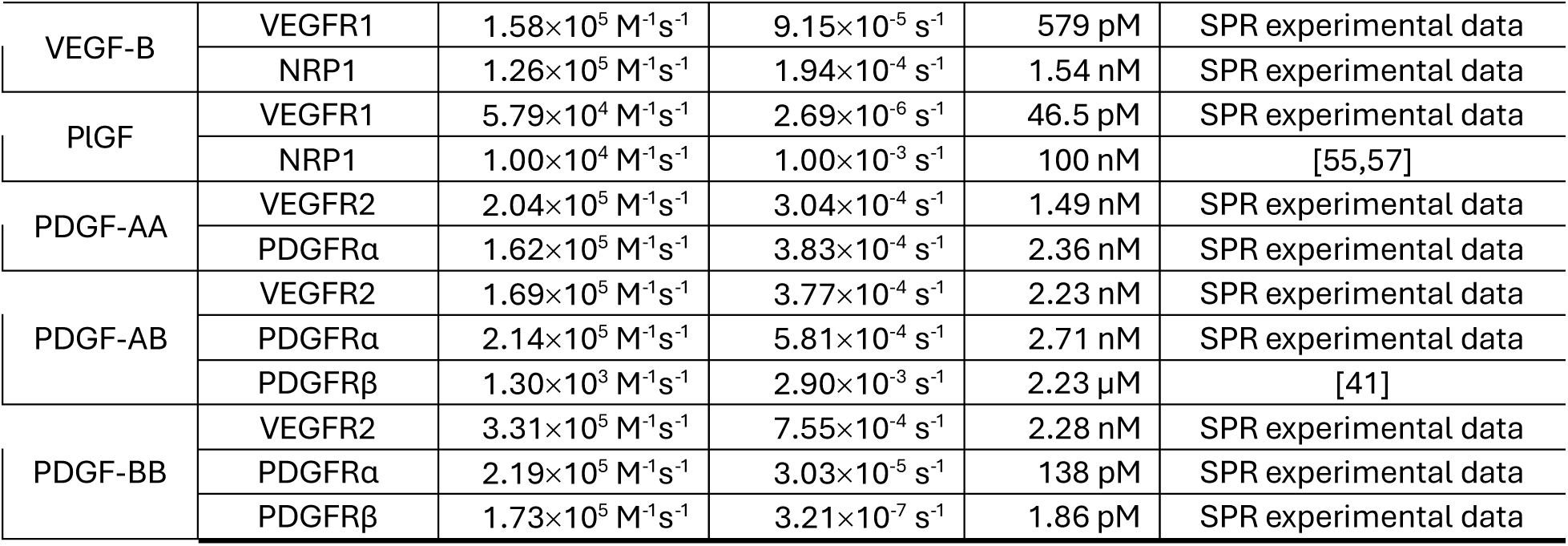
Association and dissociation rates of ligand-receptor binding.

The binding rates of ligands to receptors were obtained through literature search and surface plasmon resonance (SPR) assays conducted by Shobhan Kuila (Table 2). The detailed protocols of SPR binding assays will be stated at the end of this section. The association rate constant (k_on_) and dissociation rate constant (k_off_) for VEGF-A binding to VEGFR1, VEGFR2, and NRP1 was brought from our previous data mining work [54]. The binding affinity (K_d_) of PlGF to NRP1 was reported as 100 nM from a previous SPR study [55]. Since k_on_ and k_off_ were not described, we assumed k_off_ to be 10^-3^ s^-1^ and used the relationship K_d_ = k_off_ / k_on_ to determine k_on_ to be 10^4^ M^-1^s^-1^. This k_off_ assumption is based on assuming that PlGF:NRP1 follows VEGF dynamics as follows: 1) k_off_ = 10^-3^ s^-1^ aligns with the known VEGF-A:VEGFRs and VEGF-A:NRP1 k_off_ range: 1.00×10^-4^ to 3.50×10^-3^ s^-1^, and 2) previous studies estimated that the dissociation rate constants as 10^-3^ s^-1^ from experimental data for VEGF binding to VEGFRs on the cell surface [56].

The coupling rates of the receptors used in the model are presented in Table 3. Please note that these rates were computationally estimated by Mac Gabhhan and Popel [56]. The unit, cm^2^/mol/s, was converted to M^-1^s^-1^ by using the surface area (1.00×10^-5^ cm^2^) [58] and the volume of an endothelial cell (1 pL) [53], as mentioned above.

**Table 3.** Receptor coupling rates.

| Receptor coupling | $k_{\text{on}}$ (Measured value) | $k_{\text{on}}$ (Model value) | $k_{\text{off}}$ | Reference |
| --- | --- | --- | --- | --- |
| VEGFR1+NRP1 ↔ VEGFR1:NRP1 | $2.50 \times 10^5 \text{ M}^{-1} \text{ s}^{-1}$ | $2.50 \times 10^5 \text{ M}^{-1} \text{ s}^{-1}$ | $4.50 \times 10^{-4} \text{ s}^{-1}$ | [59] |
| VEGF-A:VEGFR2 + NRP1 ↔ VEGF-A:VEGFR2:NRP1 | $3.10 \times 10^{13} \text{ cm}^2/\text{mol/s}$ | $3.10 \times 10^6 \text{ M}^{-1} \text{ s}^{-1}$ | $1.00 \times 10^{-3} \text{ s}^{-1}$ | [56,60] |
| VEGF-A:NRP1 + VEGFR2 ↔ VEGF-A:VEGFR2:NRP1 | $1.00 \times 10^{14} \text{ cm}^2/\text{mol/s}$ | $1.00 \times 10^7 \text{ M}^{-1} \text{ s}^{-1}$ | $1.00 \times 10^{-3} \text{ s}^{-1}$ | [44,56] |

The unit “ $\text{cm}^2/\text{mol/s}$ ” was converted to $\text{M}^{-1}\text{s}^{-1}$ by multiplying $1.00 \times 10^{-7} \text{ M}/(\text{cm}^2/\text{mol/s})$ (= [Endothelial cell volume]/[Endothelial cell area]). Please note, these rates were all estimated by computational models in the citations listed, and not experimentally derived.

### Model simulation

The codes were developed to solve the ODE system with given parameter values and initial conditions for ligands and receptors above. The concentration of complexes was set to 0 at the initial time. The ‘ode15s’ and ‘pdepe’ functions were used to solve the ODE system and the PDE model in MATLAB R2024b, which will be described in the next subsection. These solvers were selected since they are appropriate for solving the stiff equations. The system of differential equations for chemical reactions could be stiff (i.e., numerically unstable if the step size is not extremely small) because of the large difference between association and dissociation rate constants. The absolute error tolerance was set to 10^-20^, and the relative error tolerance was set to 10^-9^ to control the accuracy of the solutions. The ‘NonNegative’ option was used to ensure the positivity of the solutions. The simulation was implemented in MATLAB R2024b on Mac OS X. The code is available in a GitHub repository (https://github.com/YunjeongLee/x-family_membrane).

### Model validation

All ligands concurrently participated in binding interactions. Three outcomes were determined: 1) the percentage of free and bound ligands, 2) the percentage of unoccupied and bound receptors, and 3) the percentage of each ligand occupying each receptor.

To validate our model through comparative analysis with existing frameworks, we recapitulated a computational model proposed by Mac Gabhann and Popel in 2004, which describes the *in vitro* binding of VEGF-A and PlGF to VEGFRs on endothelial cells [61]. A system of partial differential equations characterizes this model, so hereafter it will be called the “PDE model” for convenience. The parameter values used in the PDE model are presented in Table 4. Since our model and the PDE model were developed under different assumptions, it was necessary to align them within the same simulation settings. To achieve this goal, we applied one or a combination of the following conditions that were in our model to the PDE model (Fig 3): 1) matching receptor densities, 2) equal binding rates of VEGF-A and PlGF to VEGFRs, and 3) the exclusion of internalization rates for unoccupied and bound receptors, which also removes receptor synthesis. Also, since the PDE model did not consider VEGF-B, PDGF isoforms, NRP1, and PDGFRs, we set the concentrations of these species to zero in our model. While updating the conditions in the PDE model, we compared the concentrations of VEGF-A:VEGFR1 and PlGF:VEGFR1 between the PDE model and our model.

**Fig 3.**
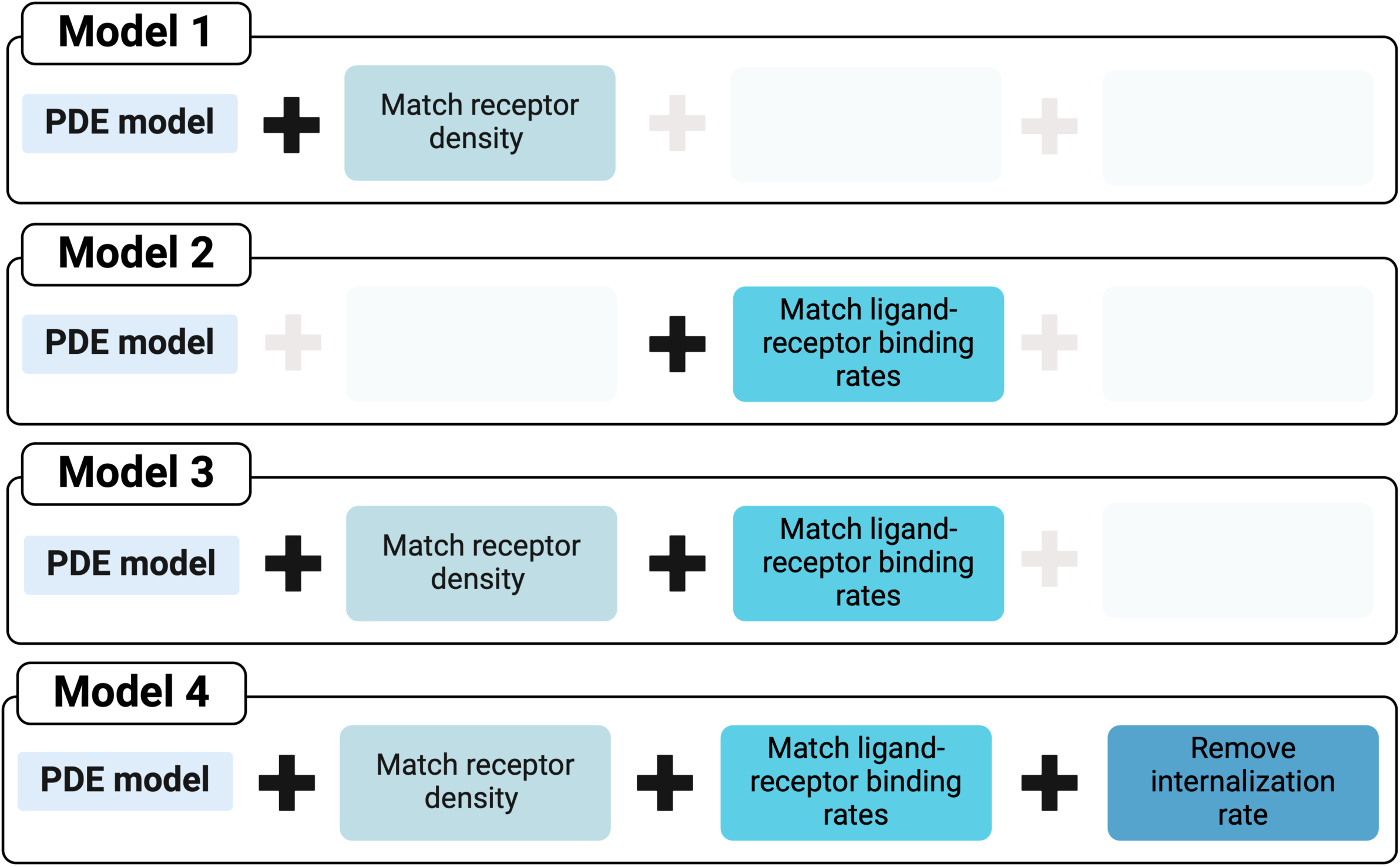
PDE Models with updated parameters. The PDE model was updated by matching receptor densities, ligand-receptor binding rates, or removing the internalization rate, which also removes the synthetic rates of receptors. In Model 1, the receptor densities in the PDE model were matched with the densities in our model. In Model 2, the binding rates of VEGF-A and PlGF to VEGFRs were matched with the rates in our model. In Model 3, both receptor densities and ligand-receptor binding rates were updated. In Model 4, the PDE model was updated by matching receptor densities and ligand-receptor binding rates, and additionally, by removing the internalization rate of receptors.

**Table 4.** Parameter values used in Mac Gabhann and Popel, 2004.

| Parameters |  | Value | Parameters |  | Value |
| --- | --- | --- | --- | --- | --- |
| Initial concentration | VEGF-A | 2.2 nM | Binding affinity | VEGF-A:VEGFR1 | 25 pM |
|  | PlGF | 2.2 nM |  | VEGF-A:VEGFR2 | 340 pM |
|  | VEGFR1 | 80,000 rec/cell |  | PlGF:VEGFR1 | 230 pM |
| | VEGFR2 | 230,000 rec/cell | Association rate constant | VEGF-A:VEGFR1 | $3.8 \times 10^6 \text{ M}^{-1} \text{ s}^{-1}$ |
| Internalization rate | Unoccupied receptor | $1 \times 10^{-5} \text{ s}^{-1}$ | | VEGF-A:VEGFR2 | $1.2 \times 10^6 \text{ M}^{-1} \text{ s}^{-1}$ |
| | Bound receptor | $2.8 \times 10^{-4} \text{ s}^{-1}$ | | PlGF:VEGFR1 | $1.5 \times 10^6 \text{ M}^{-1} \text{ s}^{-1}$ |
| Insertion rate | VEGFR1 | 0.8 rec/cell/s | Dissociation rate constant | VEGF-A:VEGFR1 | $9.5 \times 10^{-5} \text{ s}^{-1}$ |
| | VEGFR2 | 2.3 rec/cell/s | | VEGF-A:VEGFR2 | $4.1 \times 10^{-4} \text{ s}^{-1}$ |
| Diffusivity | VEGF-A | $1 \times 10^{-6} \text{ cm}^2/\text{s}$ | | PlGF:VEGFR1 | $3.45 \times 10^{-4} \text{ s}^{-1}$ |
| | PlGF | $1 \times 10^{-6} \text{ cm}^2/\text{s}$ | | | |

### Model prediction

After validating our model, to investigate the effect of upregulation or downregulation of expression of receptors, we varied the density of each receptor from 0 rec/cell to 10^5^ rec/cell and assessed its effect on five outcomes: 1) the percentage of free and bound ligands, 2) the percentage of unoccupied and bound receptors, 3) the concentration of free ligands, 4) the percentage of each ligand occupying each receptor, and 5) the concentration of each ligand-receptor complex. The selection of these outcomes came from the following four rationales: 1) Examining the concentration or percentage of free and bound ligands enhances our understanding of how cross-family interactions alter the ability of endothelial cells to capture ligands. For example, cross-family interactions may increase bound ligands by increasing the type of receptors that ligands can bind to. 2) Observing the concentration or percentage of unoccupied and bound receptors helps us understand how cross-family interactions contribute to receptor occupancy and possibilities of receptor activation. Furthermore, observing the percentage of bound receptors provides insights into the available further binding, particularly the canonical interactions that are likely to result in the most robust signaling outcome. 3) Understanding cell signaling requires preceding investigation of binding events on the cell surface. This investigation analyzes the number and the state of ligand-receptor complexes, and it facilitates the analysis of the next step in cell signaling, such as receptor activation and intracellular pathways [62]. For example, T cells started to proliferate after the number of bound interleukin-2 receptors was above a certain threshold [63]. 4) Analyzing the concentration or percentage of each ligand-receptor complex advances our understanding of the possible baseline occupancy and provides insights into the extent of the contribution of each complex to angiogenic signaling, like in the computational research of epidermal growth factor receptors [64].

### Experimental methods

#### Surface Plasmon Resonance (In-Vitro)

Surface plasmon resonance (SPR) experiments were performed at 25°C using the Reichert 4SPR system (Reichert, Inc., USA) with PEG-coated gold sensor chips containing 10% COOH (Reichert, Inc., USA #13206061). The chip was divided into four flow cells: growth factors were immobilized in channels 1 or 3, leaving channels 2 or 4 blank as references. The running buffer was 1x HBS-EP pH 7.4 (10 mM HEPES, 3 mM EDTA, 150 mM NaCl, 0.005% Tween-20). The ligand NRP1 (Cat. #3870-N1-025/CF, R&D Systems), VEGFR1 (Cat. #321-FL-050/CF, R&D Systems) and VEGFR2 (Cat. #357-KD-050/CF, R&D Systems), were immobilized using an amine coupling method. EDC (40 mg/mL) and NHS (10 mg/mL) were dissolved in water, mixed, and injected at 10 μL/min for 7 minutes to activate the surface. Proteins were diluted to 30 μg/mL in 10 mM acetate buffer (pH 4.0) and injected at 10 μL/min until the immobilization level reached ≥2000 RU, based on an R_max_ target of less than 200 RU to minimize mass transfer effects. The surface was deactivated by injecting 1M ethanolamine hydrochloride-NaOH (pH 8.5) for 7 minutes at 10 μL/min.

For kinetic analysis, analyte VEGF-B (Cat. #751-VEB-025, R&D Systems), PLGF (Cat. #264-PGB-050/CF, R&D Systems) PDGF-AA (Cat. #221-AA-050, R&D Systems), PDGF-AB (Cat. #222-AB-050, R&D Systems), and PDGF-BB (Cat. #220-BB-050, R&D Systems) were injected at concentrations of 50, 25, 12.5, 6.25, and 3.12 nM, and the association and dissociation curves were fitted using a 1:1 Langmuir binding model in TraceDrawer ver.1.8.1. Sensorgrams were visually inspected, and the fitting was validated by the χ2-to-Rmax ratio (<0.10), ensuring a reliable 1:1 interaction model. Raw sensorgrams (3.12–50 nM) were aligned, and nonspecific binding was subtracted using reference channel sensorgrams. Global fitting is considered more accurate than single curve fitting [65], so it was applied using nonlinear least squares analysis in TraceDrawer to determine association (k_on_) and dissociation (k_off_) rates across multiple response curves.

## Results

### Surface Plasmon Resonance measurement of the binding kinetics for VEGF family growth factors

Binding data for many of the growth factor interactions needed for our study were lacking throughout the literature. So, we performed SPR experiments to determine missing association and dissociation rates for missing ligand-receptor interactions: PlGF to VEGFR1 and NRP1, and VEGF-B to VEGFR1 and NRP1. We identified that PlGF binds to VEGFR1 with K_d_ of 46.5 pM (Fig S1A). We also found that VEGF-B binds predominantly to VEGFR1 with an affinity of 579 pM (Fig S1B; k_on_ = 1.58×10^5^ M^-1^s^-1^ and k_off_ = 9.15×10^-5^ s^-1^), making it three-fold stronger than its binding to NRP1, which has an affinity of 1.54 nM (Fig S1C; k_on_ = 1.26×10^5^ M^-1^s^-1^ and k_off_ = 1.94×10^-4^ s^-1^).

We determined the binding rates and affinities for the PDGF bindings to VEGFR2 (Table 2 and Figs S1D–F; PDGF-AA:VEGFR2, PDGF-AB:VEGFR2, and PDGF-BB:VEGFR2). We observed that the binding affinities of PDGF-AA to VEGFR2 was 1.49 nM with k_on_ of 2.04×10^5^ M^-1^s^-1^ and k_off_ of 3.04×10^-4^ s^-1^ (Fig S1D). The binding affinity of PDGF-AB to VEGFR2 was measured as 2.23 nM with k_on_ of 1.69×10^5^ M^-1^s^-1^ and k_off_ of 3.77×10^-4^ s^-1^ (Fig S1E), while the binding affinity of PDGF-BB to VEGFR2 was measured as 2.28 nM with k_on_ of 3.31×10^5^ M^-1^s^-1^ and k_off_ of 7.55×10^-4^ s^-1^ (Fig S1F).

We further reestablish the kinetics data of canonical interaction of PDGF bindings to PDGFRs (Table 2 and Figs S1G–J; PDGF-AA:PDGFRα, PDGF-AB:PDGFRα, PDGF-BB:PDGFRα, and PDGF-BB:PDGFRβ). PDGF-AA bound to PDGFRα with K_d_ of 2.36 nM, k_on_ of 1.62×10^5^ M^-1^s^-1^, and k_off_ of 3.83×10^-4^ s^-1^ (Fig S1G). PDGF-AB also bound to PDGFRα with the similar affinity (2.71 nM), k_on_ of 2.14×10^5^ M^-1^s^-1^, and k_off_ of 5.81×10^-4^ s^-1^ (Fig S1H). On the other hand, PDGF-BB:PDGFRα showed a stronger binding affinity (Fig S1I; K_d_ = 138 pM, k_on_ = 2.19×10^5^ M^-1^s^-1^, and k_off_ = 3.03×10^-5^ s^-1^) compared to PDGF-AA and PDGF-AB binding to PDGFRα. Finally, PDGF-BB bound to PDGFRβ with a very strong binding affinity of 1.86 pM (Fig S1J; k_on_ = 1.73×10^5^ M^-1^s^-1^ and k_off_ = 3.21×10^-7^ s^-1^).

**Fig S1.**
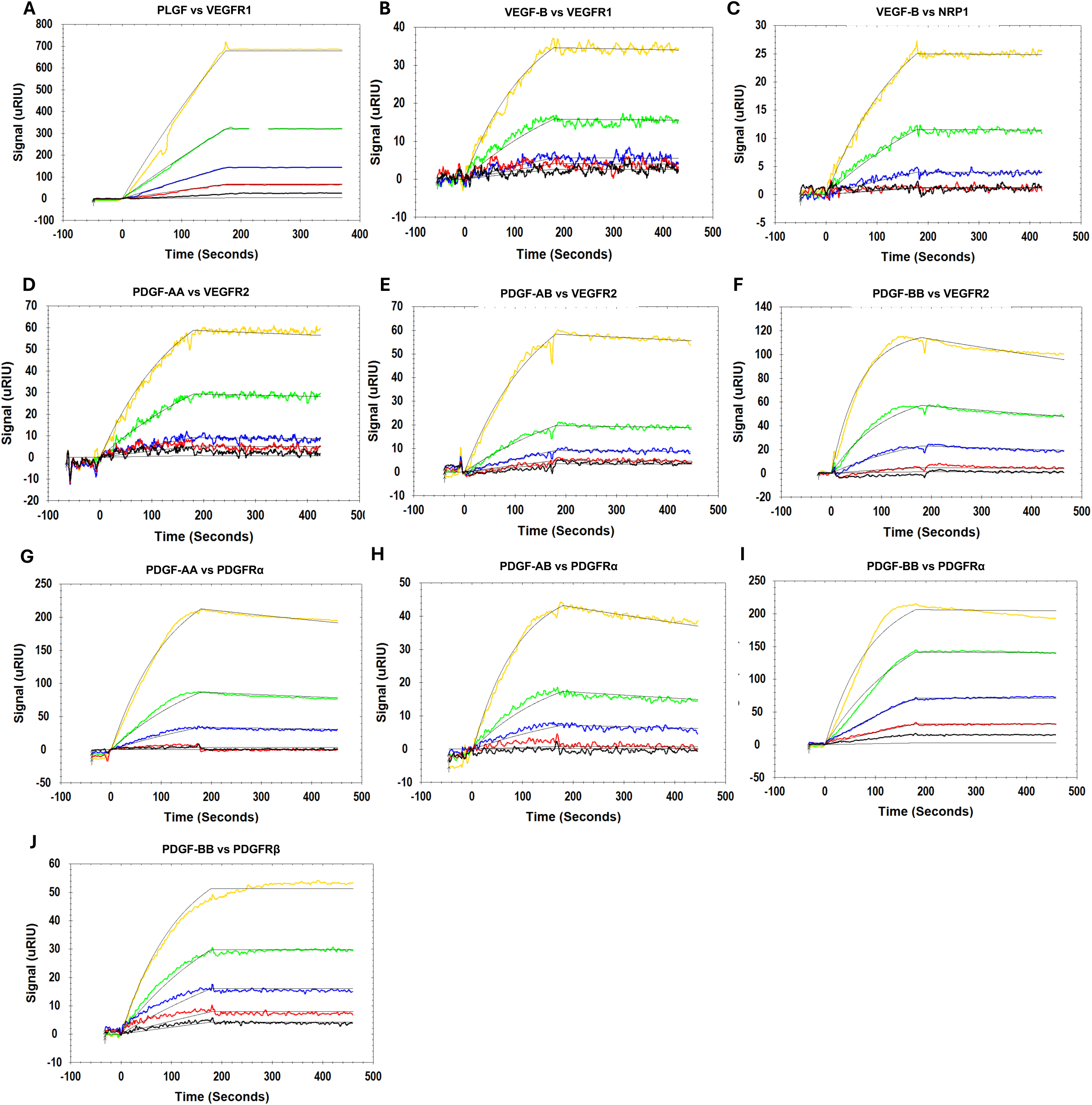
Interaction kinetics of. **(A)** PlGF vs. VEGFR1, **(B)** VEGFB vs. VEGFR1, and **(C)** VEGF-B vs. NRP1, **(D)** PDGF-AA vs. VEGFR2, **(E)** PDGF-AB vs. VEGFR2, and **(F)** PDGF-BB vs. VEGFR2, **(G)** PDGF-AA vs. PDGFRα, **(H)** PDGF-AB vs. PDGFRα, **(I)** PDGF-BB vs. PDGFRα, and **(J)** PDGF-BB vs. PDGFRβ, where VEGFR1, VEGFR2, NRP1, PDGFRα, and PDGFRβ was immobilized and analyte was passed over them at different concentrations: 50 nM (yellow), 25 nM (green), 12.5 nM (blue), 6.25 nM (red), and 3.125 nM (black). Note: The thin black overlapping lines are fitted curves of a 1:1 Langmuir model drawn with TraceDrawer ver. 1.8.1 software.

### Our model is built first by validation to the previous computational model for VEGF family interactions on endothelial cells

As an initial model validation, we recapitulated the PDE model developed by Mac Gabhann and Popel in 2004 [61] and observed similar results. More specifically, the VEGFR1 complex concentration peaked near 70,000 rec/cell (Fig S2C) and the VEGFR2 complex peaked near 160,000 rec/cell (within 17% of the prior model) (Fig S2D) with VEGF and PlGF treatment on endothelial cells. The minor discrepancy in the concentration of VEGFR2 complex (concentration of VEGFR2 complex at the peak in the PDE model in Mac Gabhann and Popel (2004): 140,000 rec/cell) may be due to the number of significant digits in the parameter values. It is noteworthy that the concentration of free VEGF and PlGF barely changed even after the ligand-receptor interactions reached the equilibrium state (Fig S2A–B). Overall, these results suggest that we successfully regenerated the VEGF and PlGF dynamics in endothelial cells in the model by Mac Gabhhann and Popel.

**Fig S2.**
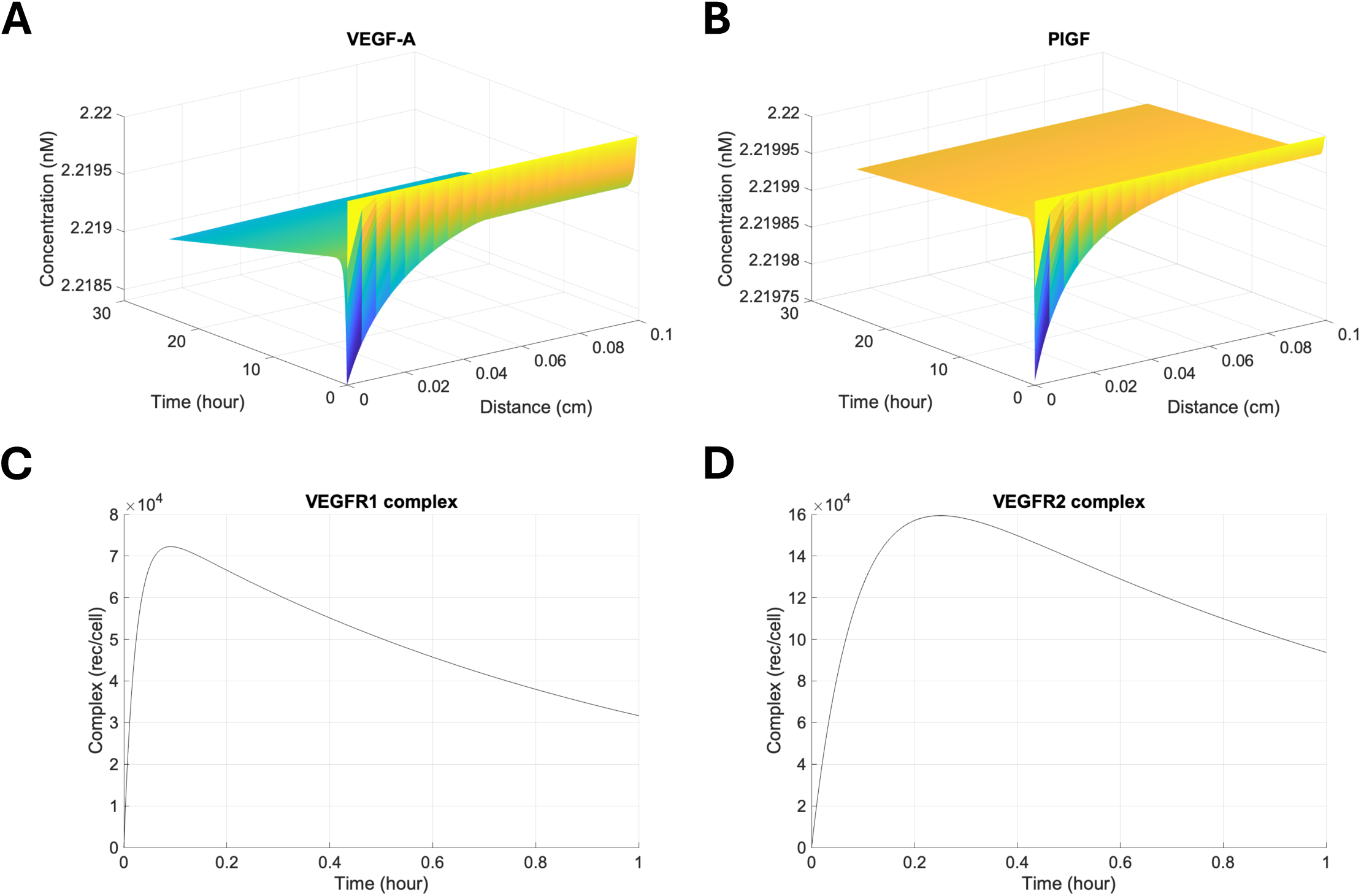
Regenerated solution for the computational model developed by Mac Gabhann and Popel in 2004. The model is a system of coupled reaction-diffusion equations, and it describes the VEGF and PlGF binding to VEGFR1 and VEGFR2 on endothelial cells in a well of a 24-well plate. The concentrations of VEGF (A) and PlGF (B) depending on space (x-axis) and time (y-axis) are presented. The “Distance” means the distance from the endothelial cell surface (i.e., *distance* = 0 cm). The concentrations of VEGFR1 (C) and VEGFR2 (D) complexes are plotted.

The difference in results between the PDE model and our model was the ratio of VEGF:VEGFR1 and PlGF:VEGFR1 concentrations. The PDE model showed VEGF-A as a dominant binding partner of VEGFR1 (Fig 2A in Mac Gabhann and Popel, 2004) while our model showed PlGF as a dominant binding partner (Fig 7). To examine if our model generates the same concentration of VEGF:VEGFR1 and PlGF:VEGFR1 as in the PDE model under the same parameter settings, we matched the settings in both models. We assumed very high concentrations of free VEGF and PlGF (2.22 µM for each) in our model to reflect the observation that the ligand concentration was maintained even after the equilibrium state was reached in the PDE model. While updating the parameter settings, we regenerated Fig 2A in Mac Gabhann and Popel, 2004, which shows the profiles of concentration of VEGFR1 complexes occupied by VEGF or PlGF for 3 hours of simulation time (Fig 4). When we updated only receptor densities in the PDE model, the concentration of VEGFR1 complexes reduced from 80,000 rec/cell to around 1,500 rec/cell (Fig 4A and 4B). However, the ratio between the concentrations of VEGF:VEGFR1 and PlGF:VEGFR1 did not change. The update in binding rates of ligands to their receptors changed the ratio whereas the total VEGFR1 complexes barely changed (Fig 4C). Updating both receptors’ densities and binding rates lowered the peak value of the VEGFR1 complexes and the proportion of PlGF:VEGFR1 among the total complexes (Fig 4D). This was because PlGF has a slower association rate with VEGFR1 compared to VEGF-A and VEGFR1 was quickly internalized after binding with VEGF-A. Thus, there would not be enough time for PlGF to bind to VEGFR1. Removal of the internalization rate, automatically removing the synthesis of new VEGFR1s and VEGFR2s in the model, led to around 10 days of the equilibrium time. This was longer than that for the previous simulations (less than 24 hours in Mac Gabhann and Popel, 2004) because of the slow dynamics of PlGF binding to receptors. The resulting percentages of VEGF and PlGF occupying VEGFR1 among the total VEGFR1 complexes at the equilibrium state were around 84% and 16%, respectively (Fig 4E). Our model also showed the same outcomes (Fig 4F) under the assumption of high concentrations of ligands (2.22 µM) compared to the receptor densities (VEGFR1: 2.66 nM and VEGFR2: 6.81 nM). Therefore, we confirmed that our model and the PDE model yielded the same results under the same model assumptions.

**Fig 4.**
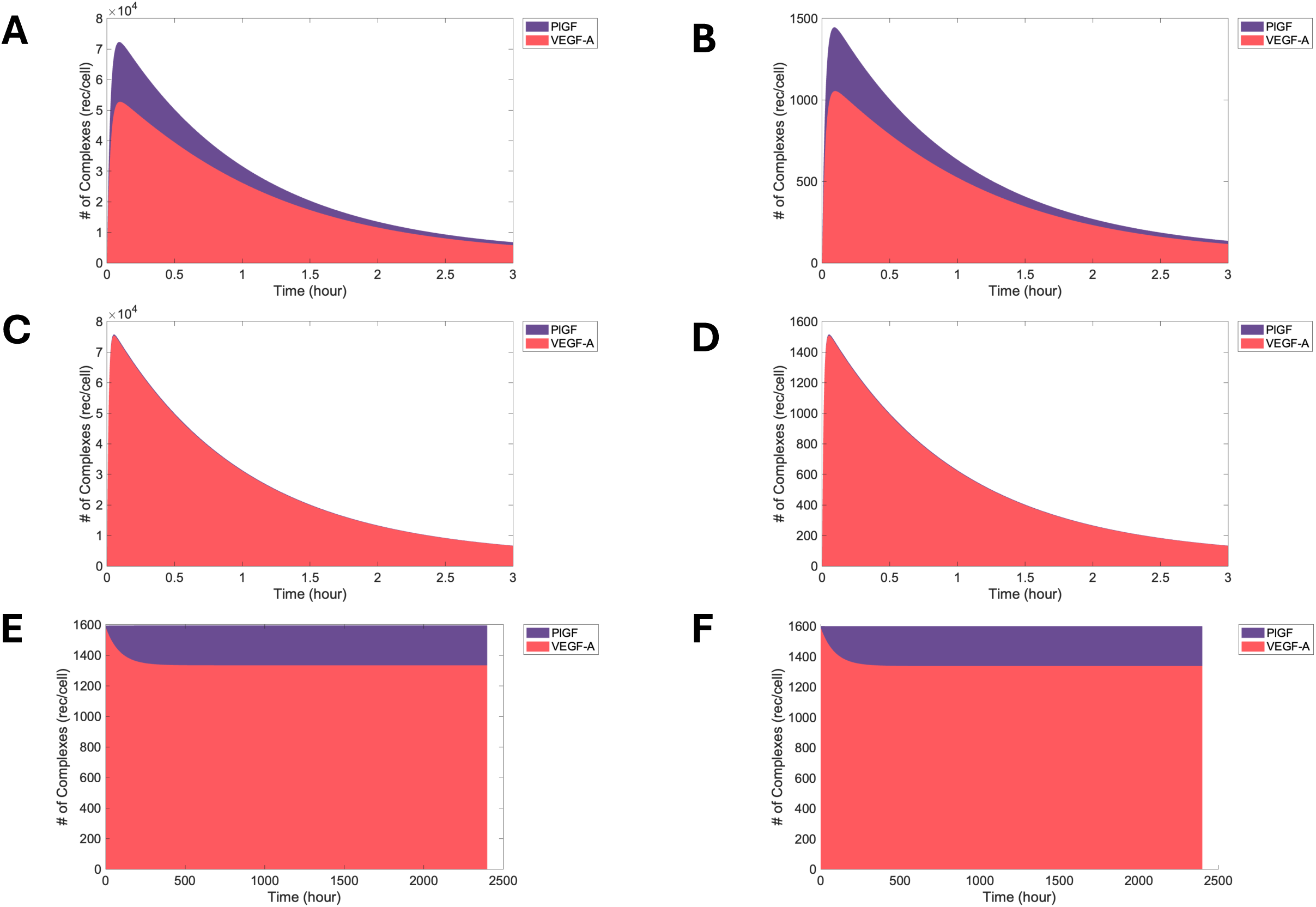
Concentrations of VEGFR1 complexes occupied by VEGF-A and PlGF predicted by the computational model developed by Mac Gabhann and Popel, under various assumptions and our model. The concentrations of VEGF-A:VEGFR1 and PlGF:VEGFR1 predicted from the PDE model are represented by red and purple areas when (A) the same parameters in Mac Gabhann and Popel, 2004 was used, (B) the densities of VEGFR1 and VEGFR2 are updated, (C) the binding rates of VEGF-A and PlGF to VEGFRs are updated, (D) both the receptor densities and binding rates are updated, and (E) both the receptor densities and binding rates are updated, and no internalization of free and bound receptors is assumed. (F) With the high concentration of free ligands in our model, the proportion of VEGF-A and PlGF bound to VEGFR1 is around 84% and 16%, respectively, among the total VEGFR1 complexes.

### Long equilibrium time due to slow PlGF dynamics

After the model validation, we set the concentration of all ligands as 1 nM. This ligand concentration was chosen since it is commonly used to stimulate endothelial cells and it is considered to be optimal in the cell signaling [46–50]. The receptor densities were also restored so that our model included VEGFR1, VEGFR2, NRP1, PDGFRα, and PDGFRβ on endothelial cells. Our model showed that the equilibrium state was reached ten days after the initiation of interactions between ligands and receptors (Fig 5). With the exception of PlGF, the ligands rapidly bound to their receptors within a timeframe of 20 minutes. The slower dynamics observed with PlGF were attributed to its binding affinity to NRP1, which is 37–65 times weaker compared to the other ligands, and the slow dissociation rate of PlGF:VEGFR1.

**Fig 5.**
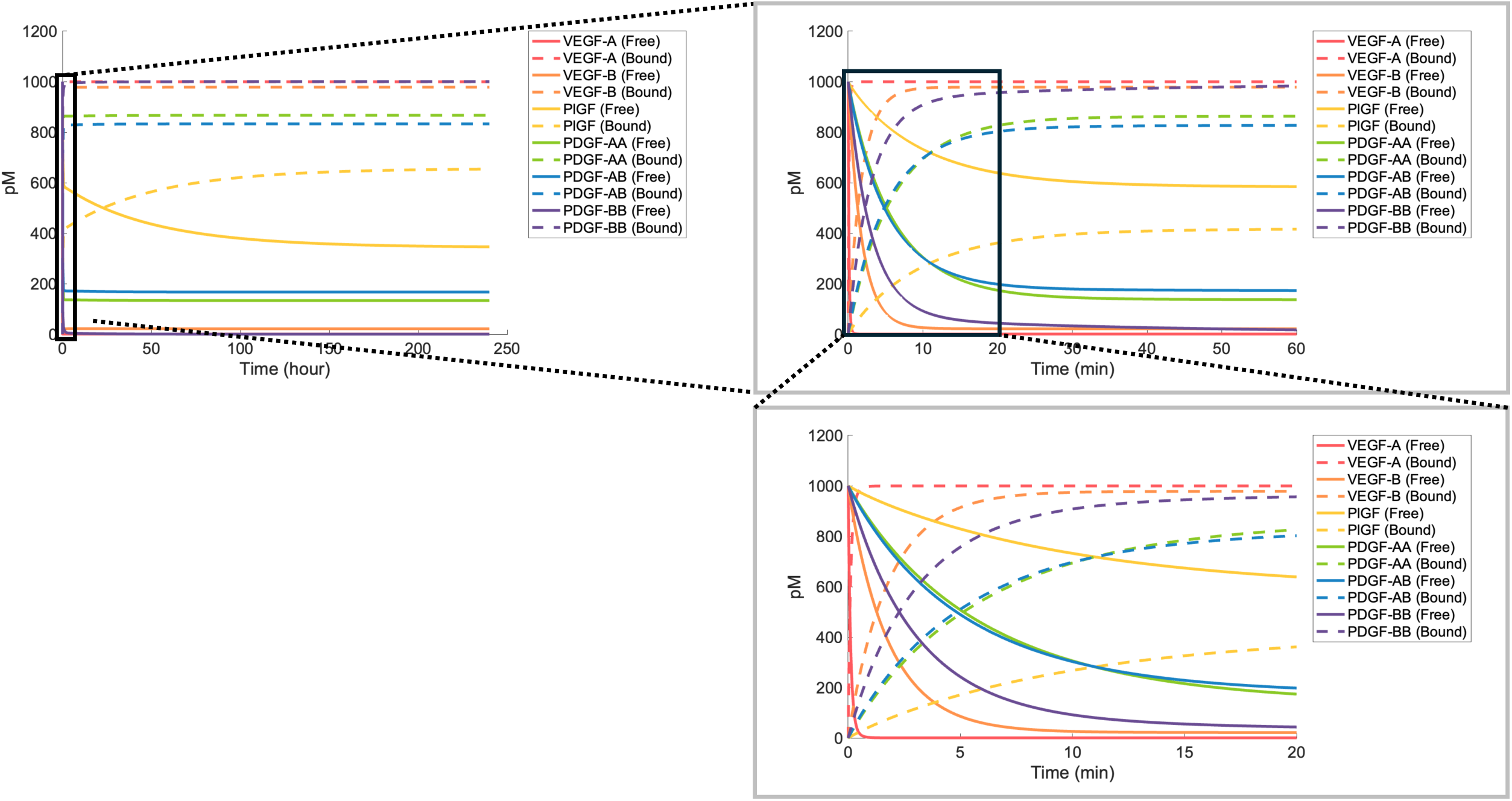
Dynamics of free and bound ligands for ten days. The concentration of ligands, i.e., VEGF-A, VEGF-B, PlGF, PDGF-AA, PDGF-AB, and PDGF-BB, are represented by colors in the order of red, orange, yellow, green, blue, and purple. The concentration of free ligands (solid line) and the bound ligands (dashed line) reached the equilibrium state ten days after treatment of all ligands. The left figure shows the dynamics of ten days after the treatment, the upper right figure shows the first one-hour’s dynamics after the treatment, and the bottom right figure shows the dynamics of the first 20 minutes after the treatment.

### All VEGF-A and PDGF-BB are bound to receptors while 3–20% of receptors are occupied

Our model showed that nearly 100% of VEGF-A and PDGF-BB were in the bound state (Fig 6A). The exceedingly high percentage of bound VEGF-A may be attributed to its ability to bind to all five receptors. The high percentage of bound PDGF-BB may be attributed to the strong binding affinity for PDGF-BB:PDGFRβ complex. Among other VEGF family members, 98% of VEGF-B was bound to their receptors, whereas 65% of PlGF was bound (Fig 6A). On the other hand, 87% of PDGF-AA and 83% of PDGF-AB were bound to VEGFR2 or PDGFRs. Additionally, the simulation revealed that 16% of VEGFR1, 26% of VEGFR2, and around 11% of PDGFRs were bound to their ligands (Fig 6B). On the other hand, only 3% of NRP1 was bound to ligands. This was due to the higher density of NRP1 (44,090 rec/cell; 73 nM) compared to the concentration of ligands (1 nM for each ligand).

**Fig 6.**
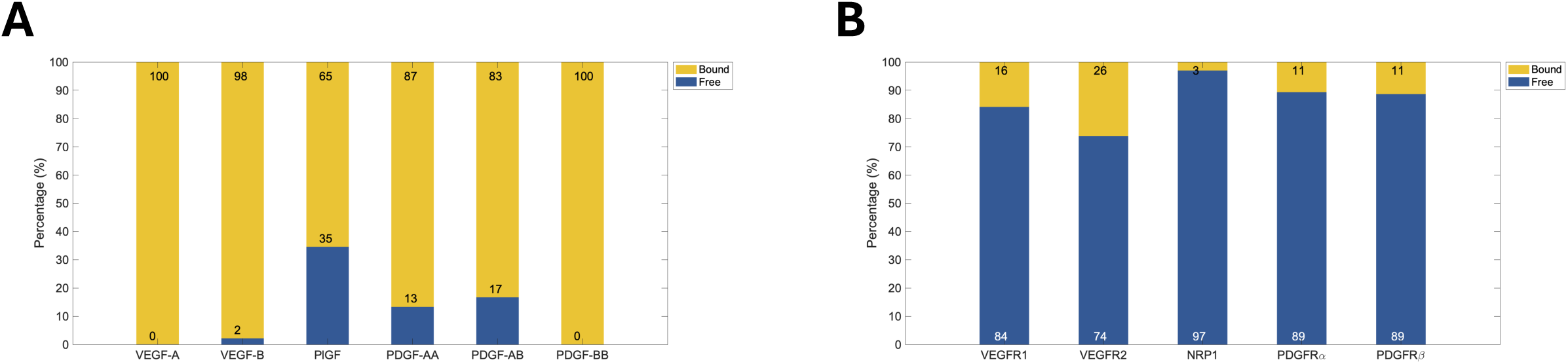
The percentage of free and bound species at the equilibrium state. The percentages of free and bound ligands (A) and the percentages of unoccupied and bound receptors (B) are shown. The yellow and orange colors represent bound and free species, respectively.

**Fig 7.**
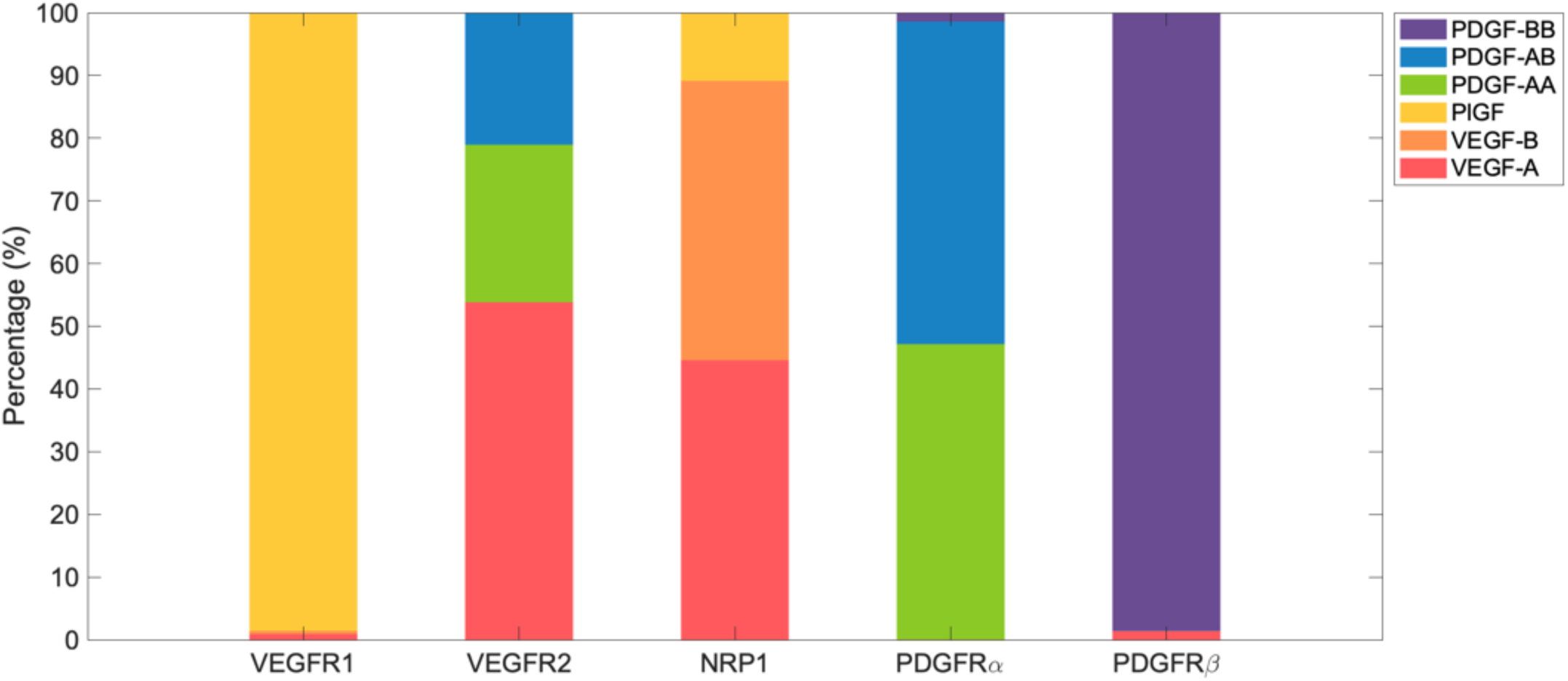
The ligand distribution on each receptor at the equilibrium state. The percentages of ligands on each receptor are represented by colors; VEGF-A (red), VEGF-B (orange), PlGF (yellow), PDGF-AA (green), PDGF-AB (blue), PDGF-BB (purple).

### Most VEGFR1 is bound to PlGF rather than VEGF-A, and VEGFR2 is bound to VEGF-A, PDGF-AA, or PDGF-AB

Identifying which ligand is bound to a receptor is important because the role of each complex is different. We thus examined the percentage of ligands that occupy each receptor (Fig 7). Most VEGFR1s were occupied by PlGF at the equilibrium state, even though VEGF-A:VEGFR1 binding is five times stronger than PlGF:VEGFR1 (9.09 pM vs. 46.5 pM). Around 50% of bound VEGFR2 was occupied by VEGF-A, and the other 50% of VEGFR2 was occupied by PDGF-AA and PDGF-AB. Nearly all PDGF-BB were bound to PDGFRβ — over 95% of PDGFRβ was bound to PDGF-BB, and the remaining PDGFRβ was occupied by VEGF-A. Almost over 95% of bound PDGFRα was bound to PDGF-AA or PDGF-AB. The remaining bound PDGFRα was in the form of PDGF-BB:PDGFRα. Finally, around 45% NRP1 was occupied by VEGF-A, 45% to VEGF-B, and the other 10% was occupied by PlGF.

### Change of VEGFR1 density depletes the concentration of free PlGF

Receptor densities have previously been established to significantly affect complex dynamics [66–68]. VEGFR1 density is a crucial regulator of ligand-receptor complex formation, particularly for PlGF, VEGF-A, and VEGF-B and their binding partners. Our model predicted that VEGFR1 density drove a shift in ligand binding preference toward VEGFR1, depleting the pool of free PlGF, and altering the distribution of VEGF family members among VEGFRs and NRP1. In contrast, PDGFR-mediated signaling remained unaffected by VEGFR1 levels. We achieved these predictions by increasing VEGFR1 density on the cell surface from 0 rec/cell to 10^5^ rec/cell and measuring five outcomes: 1) the percentage of free and bound ligands, 2) the concentration of free ligands, 3) the percentage of unoccupied and bound receptors, 4) the percentage of ligands occupying each type of bound receptors, and 5) the concentration of ligands bound to each receptor type.

As VEGFR1 density increased, the concentration of free PlGF dramatically decreased due to the preferential formation of PlGF:VEGFR1 complexes while other growth factors showed robust concentrations (Figs 8A–F and 9A–F). PlGF preferentially bound to VEGFR1 rather than NRP1 (Fig 12C) because of the lower dissociation rate of PlGF:VEGFR1 compared with PlGF:NRP1 (2.69×10^-6^ s^-1^ for VEGFR1 and 10^-3^ s^-1^ for NRP1). When the VEGFR1 density was 0 rec/cell, around 60% of PlGF was in the free state, the remaining PlGF being bound to NRP1 (Fig 8C and 12C). The depletion of PlGF from NRP1 binding sites shifted NRP1 binding primarily to its other partners, VEGF-A and VEGF-B at high VEGFR1 densities (Fig 11C).

**Fig 8.**
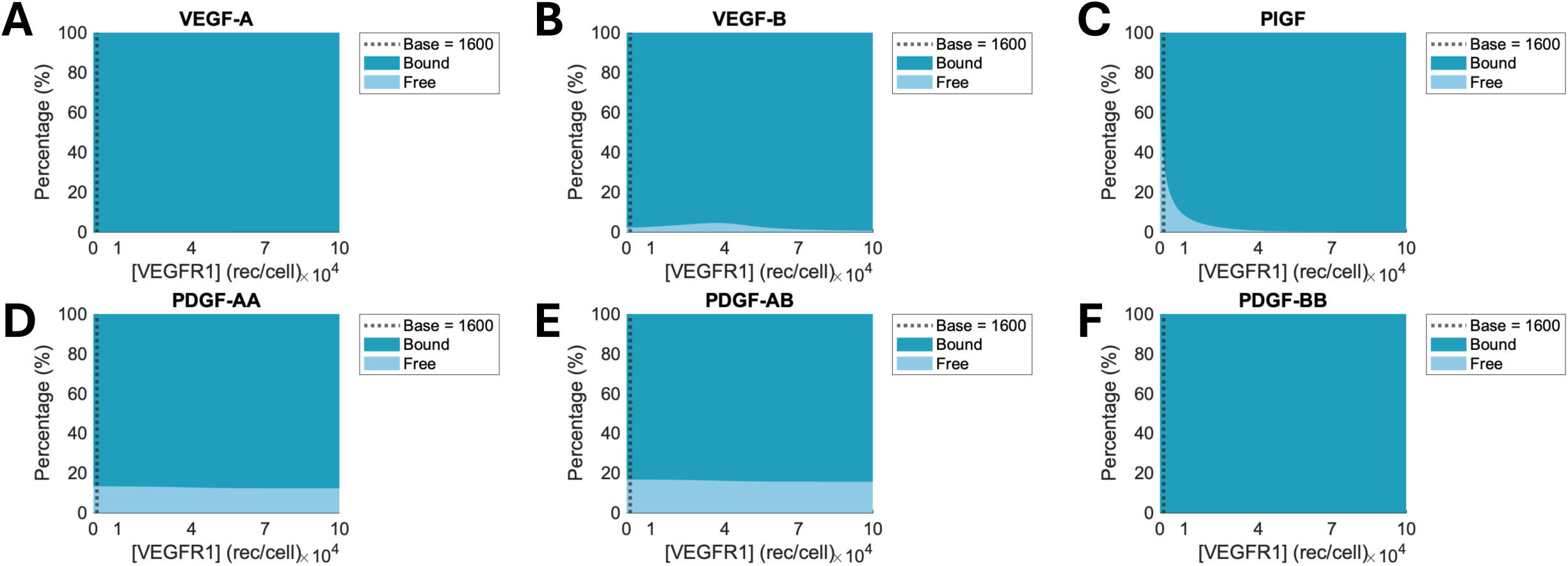
The percentage of free and bound ligands at the equilibrium state depending on the various VEGFR1 densities. The percentages of the concentrations of free and bound VEGF-A (A), VEGF-B, (B), PlGF (C), PDGF-AA (D), PDGF-AB (E), and PDGF-BB (F) at each VEGFR1 density are plotted. The light blue area represents the percentage of free ligands, while the dark blue area represents the percentage of bound ligands for each receptor density. The baseline of VEGFR1 density is 1,600 rec/cell.

**Fig 9.**
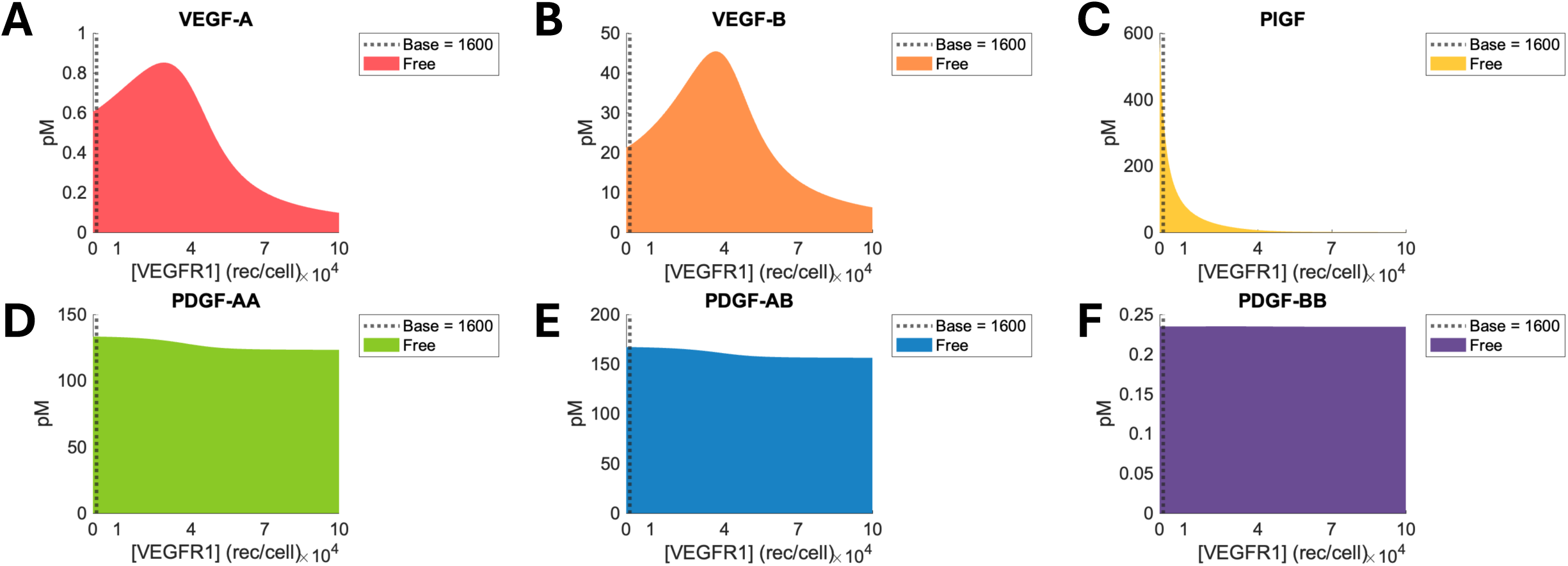
The concentration of free ligands at the equilibrium state depending on the various VEGFR1 densities. The colored area represents the concentration of free VEGF-A (A), VEGF-B (B), PlGF (C), PDGF-AA (D), PDGF-AB (E), and PDGF-BB (F) for each density of the receptor. The baseline of VEGFR1 density is 1,600 rec/cell.

**Fig 10.**
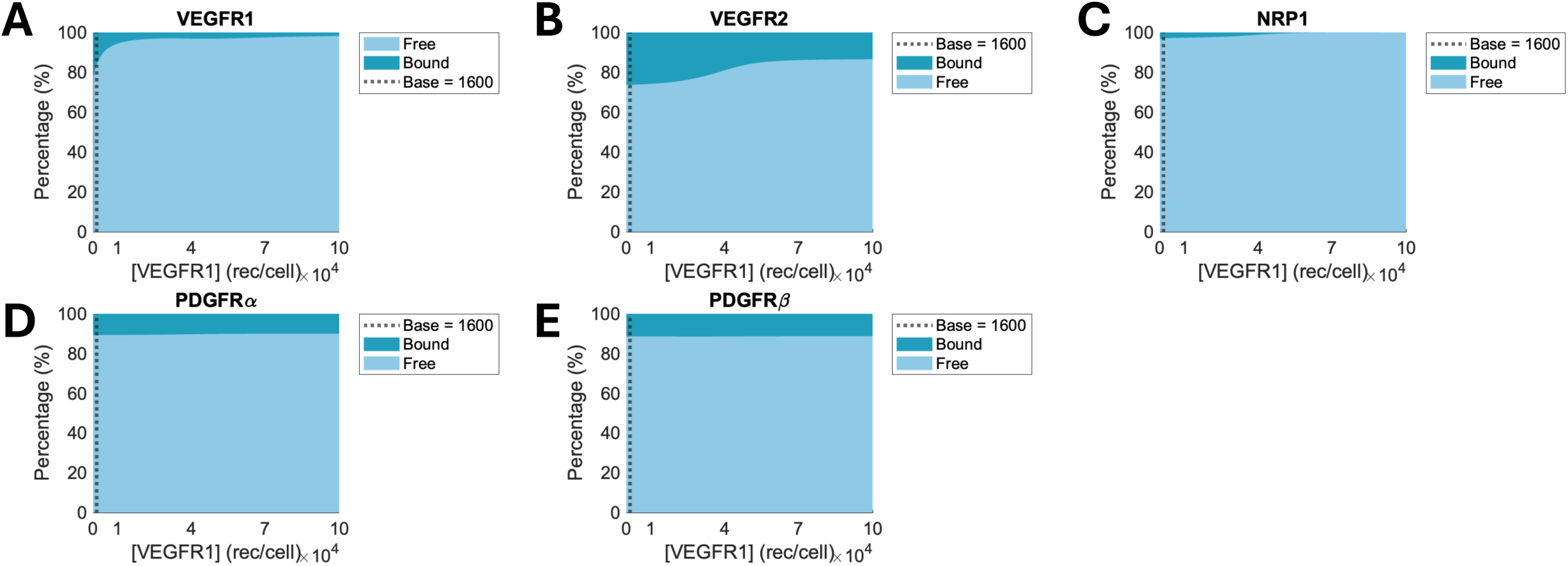
The percentage of free and bound receptors on the endothelial cell at the equilibrium state depending on the various VEGFR1 densities. The percentages of free and bound VEGFR1 (A), VEGFR2 (B), NRP1 (C), PDGFRα (D), and PDGFRβ (E) are represented. The light blue area represents the percentage of free receptors, while the dark blue area represents the percentage of bound receptors for each receptor density. The baseline of VEGFR1 density is 1,600 rec/cell.

**Fig 11.**
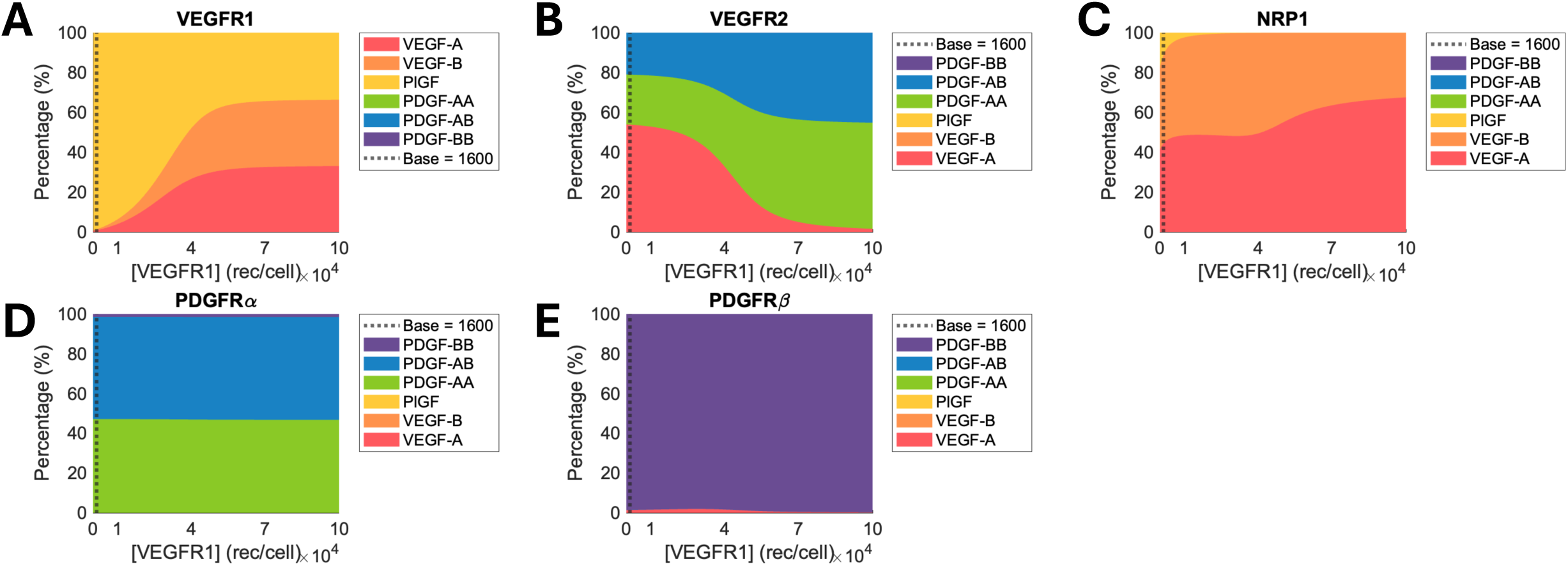
The percentage of ligands occupying each type of bound receptor at the equilibrium state depending on the various VEGFR1 densities. The percentages of VEGF-A, VEGF-B, PlGF, PDGF-AA, PDGF-AB, and PDGF-BB bound to VEGFR1 (A), VEGFR2 (B), NRP1 (C), PDGFRα (D), and PDGFRβ (E) are represented. The baseline value of the VEGFR1 density is 1,600 rec/cell.

**Fig 12.**
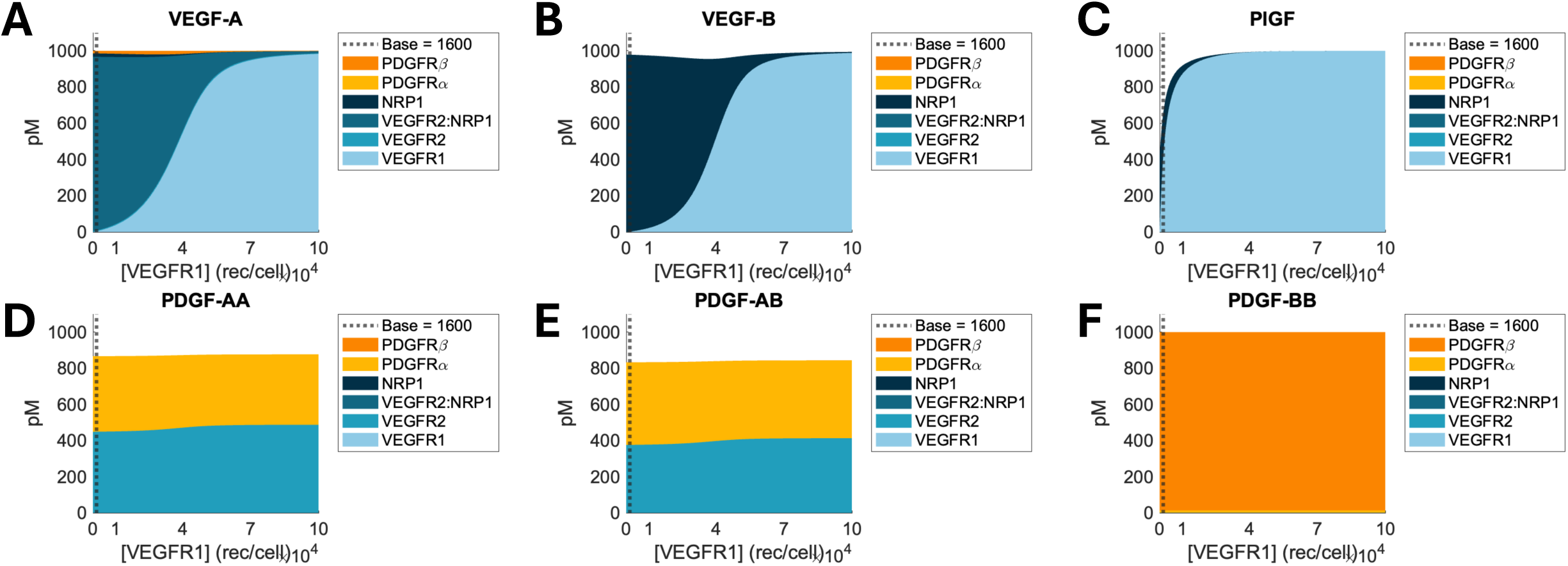
The concentration of ligand–receptor complex at the equilibrium state depending on the various VEGFR1 densities. The concentrations of VEGFR1, VEGFR2, NRP1, PDGFRα, and PDGFRβ that are bound to VEGF-A (A), VEGF-B (B), PlGF (C), PDGF-AA (D), PDGF-AB (E), and PDGF-BB (F) are represented. The baseline value of the VEGFR1 density is 1,600 rec/cell.

As VEGFR1 density increased, VEGF-A and VEGF-B also predominantly bound to VEGFR1, significantly reducing their binding to VEGFR2 and NRP1 (Figs 12A–B). The percentage of VEGF-A:VEGFR2 and VEGF-A:VEGFR2:NRP1 complexes decreased as VEGFR1 density increased, shifting the VEGF-A binding preference from VEGFR2 to VEGFR1 (Fig 12A). This resulted in a decreased percentage of bound VEGFR2 (Fig 10B) and an increased percentage of PDGFs:VEGFR2 (Fig 11B) because, contrary to the reduction in VEGF-A:VEGFR2 complexes (Fig 12A), the number of PDGF-AA:VEGFR2 and PDGF-AB:VEGFR2 complexes showed very slight increments by 30–40 pM (Fig 12D–E). The increase in VEGF-A:VEGFR1 and VEGF-B:VEGFR1 complexes was much greater than the PlGF:VEGFR1 formation (Fig 12A–C), leading to an increment of their percentages among the total bound VEGFR1s (Fig 11A). VEGF-B initially bound to NRP1 but switched to VEGFR1 as VEGFR1 density rose (Fig 12B). As a result, NRP1 showed some reduction in free binding capacity, but the effect was relatively small (3% decrease) compared to VEGFR2 due to its high baseline density (Fig 10C). When the VEGFR1 density reached 10^5^ rec/cell, almost all VEGF-A, VEGF-B, and PlGF were bound to VEGFR1 (Fig 12A–C), leading to comparable percentages between the complexes (Fig 11A). The percentage of unoccupied VEGFR1 reduced as the VEGFR1 density increased due to limited complex formation compared to increasing VEGFR1 density (Fig 10A). Unoccupied PDGFRs were unaffected by varied VEGFR1 density because their main binding partners were PDGFs, which do not bind to VEGFR1 (Fig 10D–E, 11D–E, and 12D–F).

### Increased VEGFR2 density leads to decreased PDGF:PDGFRα complexes

VEGFR2 is a main driver in VEGF-A-induced angiogenesis. Our model predicted that increasing VEGFR2 density shifted the binding preferences of PDGF-AA and PDGF-AB from PDGFRα to VEGFR2, with a reduction in free PDGFs and an increase in unoccupied PDGFRα.

As VEGFR2 density increased, free PDGF-AA and PDGF-AB reduced (Figs 13D–E and 14D–E) due to an increase in their binding to VEGFR2 (Fig 17D–E). The increased PDGFs:VEGFR2 binding altered the ligand occupancy in VEGFR2 by shifting the main binding partner of VEGFR2 from VEGF-A to PDGFs (Fig 16B). The subsequent, reduced PDGF-AA:PDGFRα and PDGF-AB:PDGFRα binding (Fig 17D–E) resulted in reduced bound PDGFRα (Fig 15D) and occupation of PDGF-AA and –AB in PDGFRα (Fig 16D). Free VEGF-A was depleted due to increased VEGFR2 density (Figs 13A and 14A), while free PDGF-BB concentration remained unaffected (Figs 13F and 14F). This was because PDGF-BB strongly binds to PDGFRβ, thus, PDGF-BB:PDGFRβ formation was the major form of PDGF-BB regardless of VEGFR2 density (Figs 16E and 17F). This robust PDGF-BB:PDGFRβ formation led to stable unoccupied PDGFRβ percentage across varying VEGFR2 density (Fig 15E).

**Fig 13.**
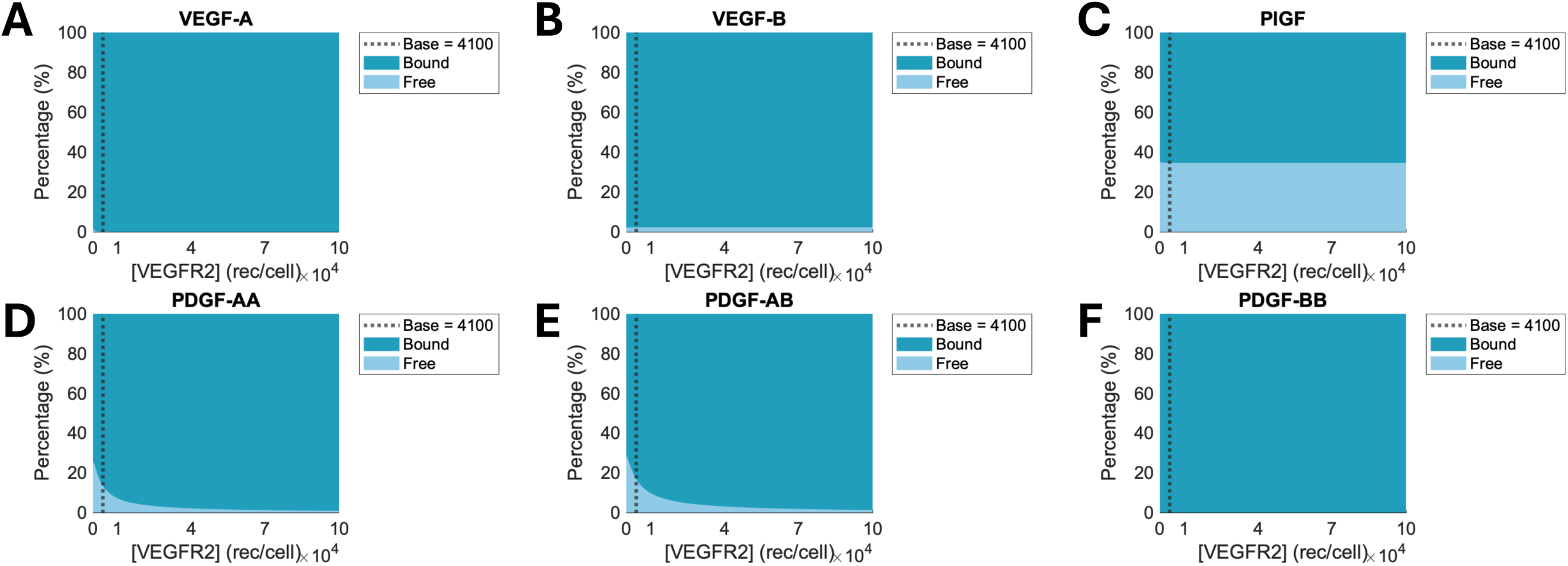
The percentage of free and bound ligands at the equilibrium state depending on the various VEGFR2 densities. The percentages of the concentrations of free and bound VEGF-A (A), VEGF-B, (B), PlGF (C), PDGF-AA (D), PDGF-AB (E), and PDGF-BB (F) at each VEGFR2 density are plotted. The light blue area represents the percentage of free ligands, while the dark blue area represents the percentage of bound ligands for each receptor density. The baseline of VEGFR2 density is 4,100 rec/cell.

**Fig 14.**
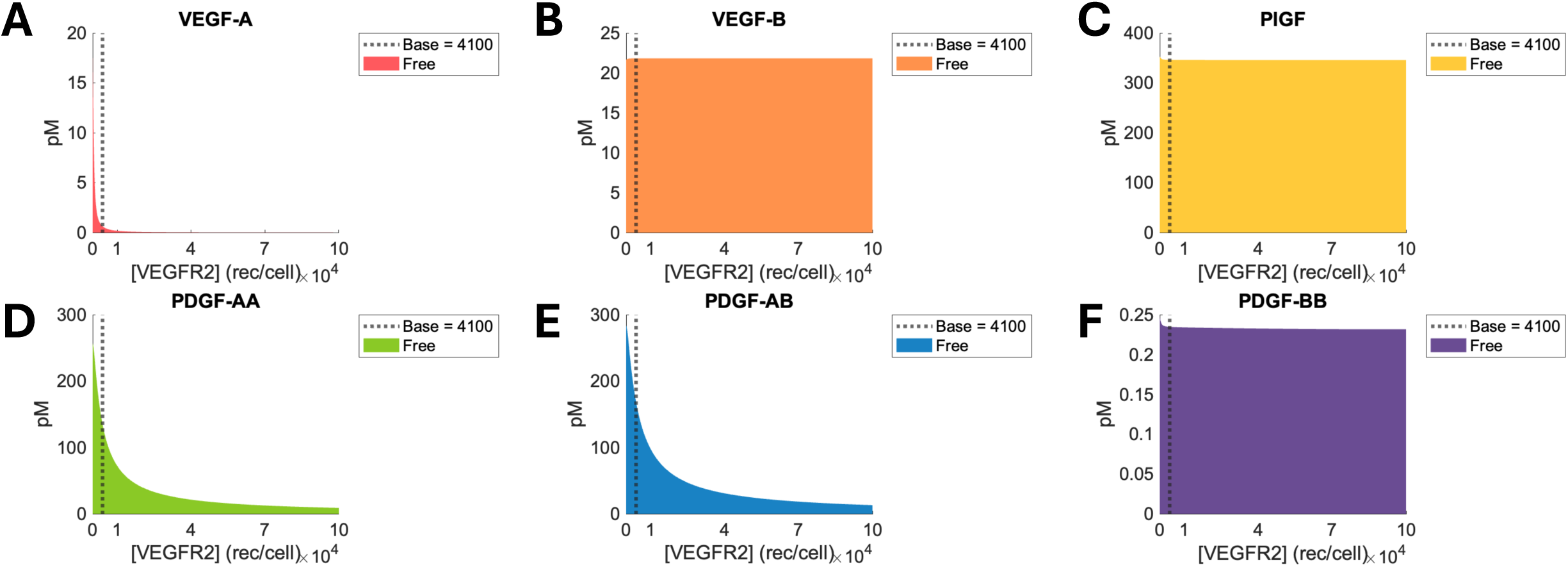
The concentration of free ligands at the equilibrium state depending on the various VEGFR2 densities. The colored area represents the concentration of free VEGF-A (A), VEGF-B (B), PlGF (C), PDGF-AA (D), PDGF-AB (E), and PDGF-BB (F) for each density of the receptor. The baseline of VEGFR2 density is 4,100 rec/cell.

**Figure 15.**
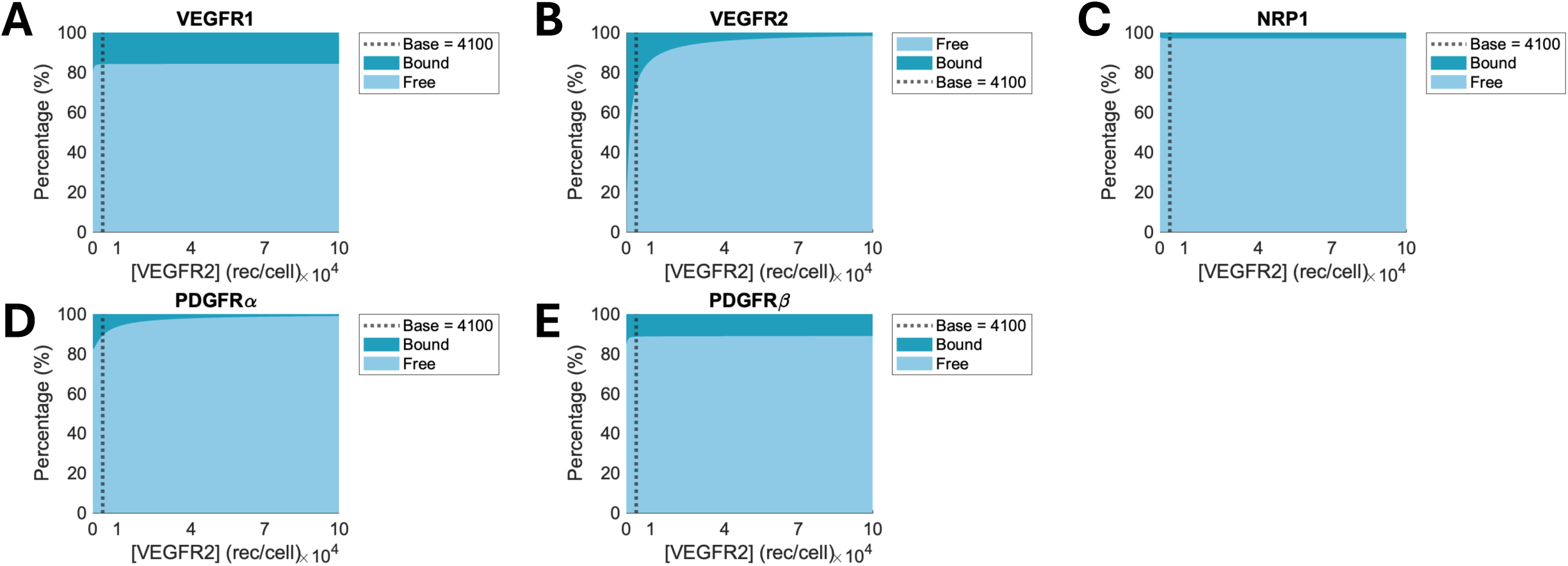
The percentage of free and bound receptors on the endothelial cell at the equilibrium state depending on the various VEGFR2 densities. The percentages of free and bound VEGFR1 (A), VEGFR2 (B), NRP1 (C), PDGFRα (D), and PDGFRβ (E) are represented. The light blue area represents the percentage of free receptors, while the dark blue area represents the percentage of bound receptors for each receptor density. The baseline of VEGFR2 density is 4,100 rec/cell.

**Fig 16.**
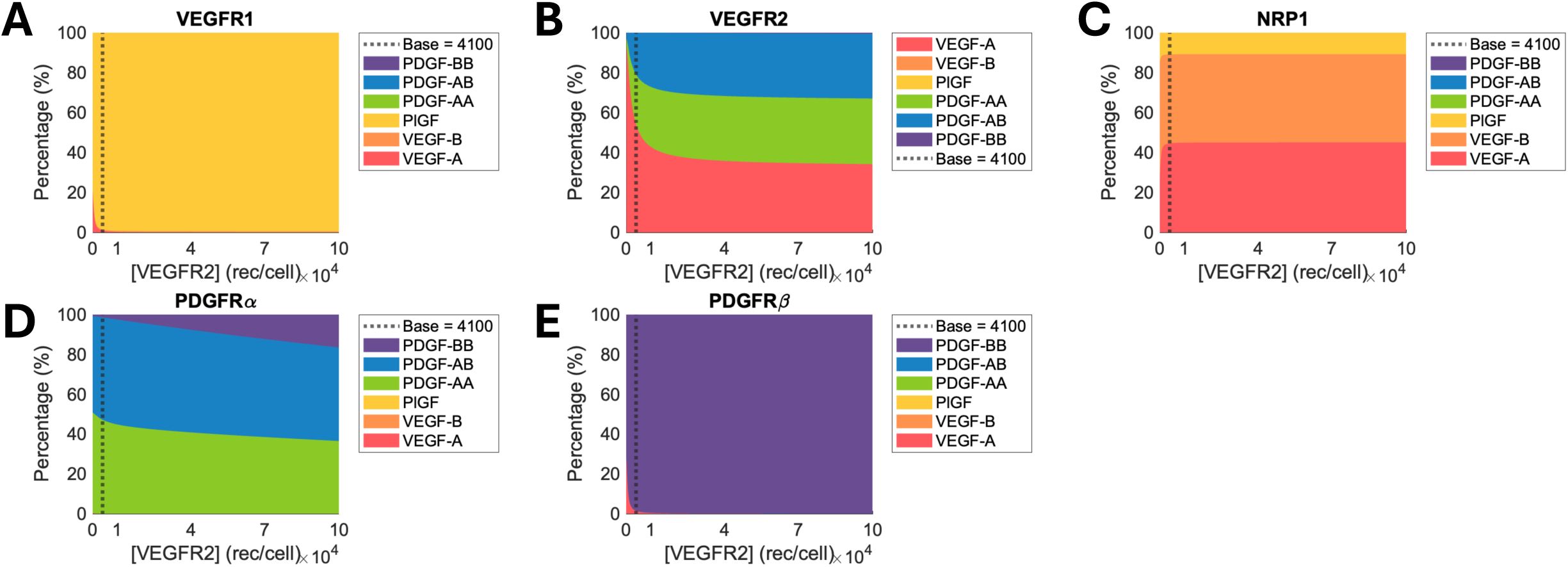
The percentage of ligands occupying each type of bound receptor at the equilibrium state depending on the various VEGFR2 densities. The percentages of VEGF-A, VEGF-B, PlGF, PDGF-AA, PDGF-AB, and PDGF-BB bound to VEGFR1 (A), VEGFR2 (B), NRP1 (C), PDGFRα (D), and PDGFRβ (E) are represented. The baseline value of the VEGFR2 density is 4,100 rec/cell.

**Fig 17.**
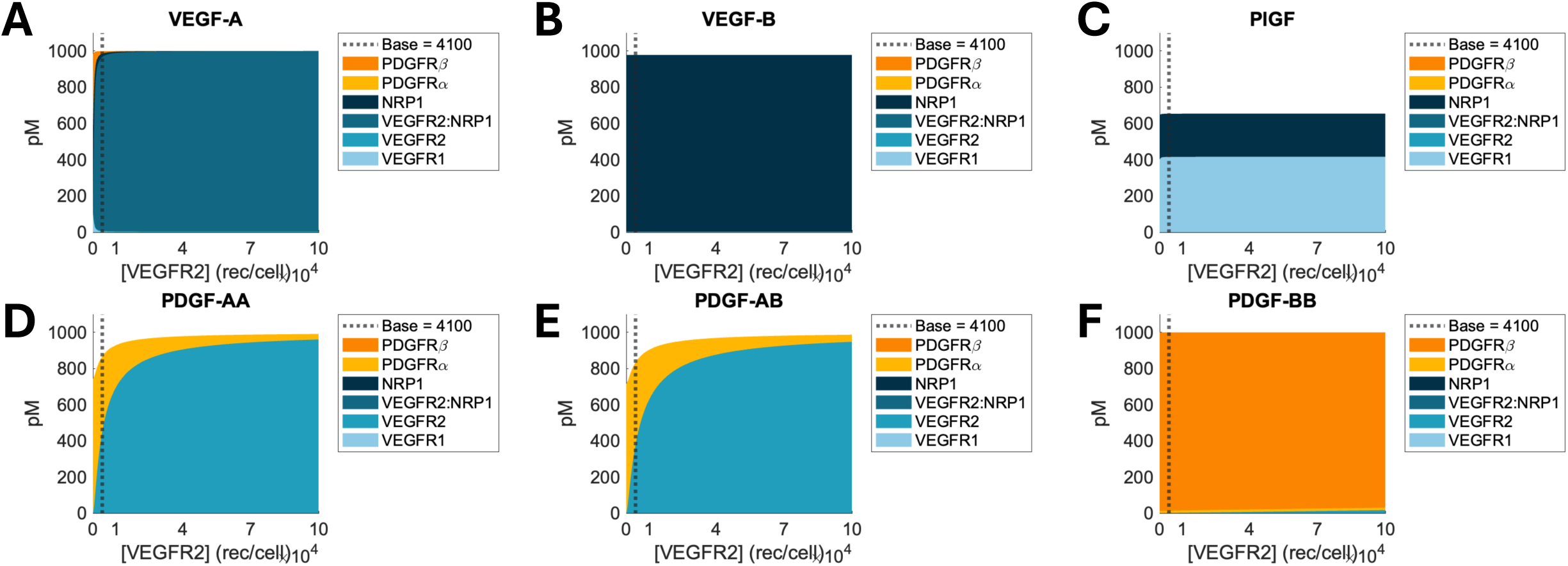
The concentration of ligand–receptor complex at the equilibrium state depending on the various VEGFR2 densities. The concentrations of VEGFR1, VEGFR2, NRP1, PDGFRα, and PDGFRβ that are bound to VEGF-A (A), VEGF-B (B), PlGF (C), PDGF-AA (D), PDGF-AB (E), and PDGF-BB (F) are represented. The baseline value of the VEGFR2 density is 4,100 rec/cell.

Increasing VEGFR2 density did affect neither the percentages of unoccupied VEGFR1 and NRP1 nor their ligand occupancy (Figs 15A, 15C, 16A, and 16C). This was for two reasons: 1) their binding partner, VEGF-B and PlGF, were unaffected by VEGFR2 density (Figs 13B–C, 14B–C, and 17B–C) because they do not bind to VEGFR2 and 2) most VEGF-A formed VEGF-A:VEGFR2 or VEGF-A:VEGFR2:NRP1 complexes regardless of VEGFR2 density (Fig 17A). Finally, increased VEGFR2 density decreased the percentage of unoccupied VEGFR2 because of limited complex formation compared to increasing its total density (Fig 15B).

### Increased NRP1 density leads to a biphasic concentration of free PlGF

NRP1 not only plays a role as a co-receptor in VEGF-A:VEGFR2 binding but also binds to VEGF-B and PlGF. Our model found that free PlGF concentration showed biphasic patterns as NRP1 density increased. When NRP1 density was increased from 0 rec/cell to the baseline, free PlGF concentration increased (Figs 18C and 19C). This was due to the faster dissociation of PlGF:NRP1 than PlGF:VEGFR1. When the NRP1 density was 0 rec/cell, PlGF bound to VEGFR1, however, as the NRP1 density increased, the formation of PlGF:NRP1 increased (Fig 22C). The fast dissociation of PlGF from NRP1 and their low binding affinity could not compensate for the loss of PlGF:VEGFR1. When NRP1 density increased from baseline to 10^5^ rec/cell, the increased PlGF:NRP1 formation reduced the free PlGF concentration (Figs 18C, 19C, and 22C). This contributed to the increase in occupancy of PlGF in bound NRP1 (Fig 21C).

**Fig 18.**
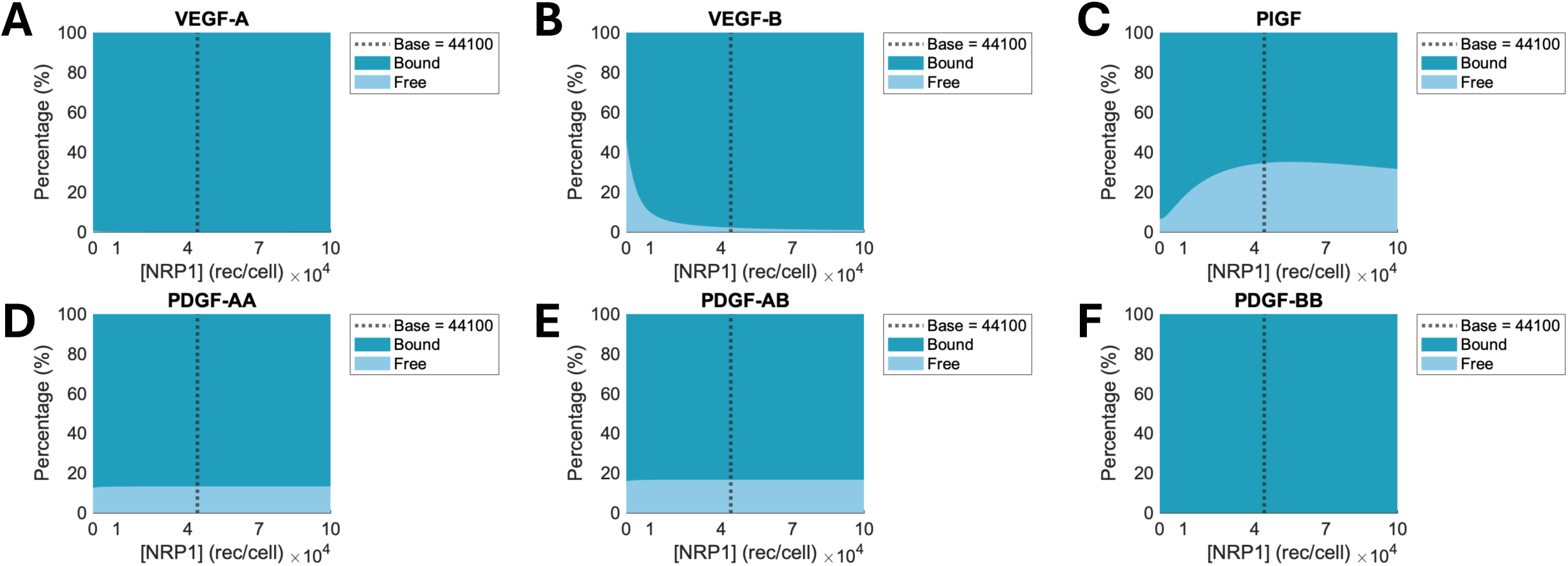
The percentage of free and bound ligands at the equilibrium state depending on the various NRP1 densities. The percentages of the concentrations of free and bound VEGF-A (A), VEGF-B, (B), PlGF (C), PDGF-AA (D), PDGF-AB (E), and PDGF-BB (F) at each NRP1 density are plotted. The light blue area represents the percentage of free ligands, while the dark blue area represents the percentage of bound ligands for each receptor density. The baseline of NRP1 density is 44,100 rec/cell.

**Fig 19.**
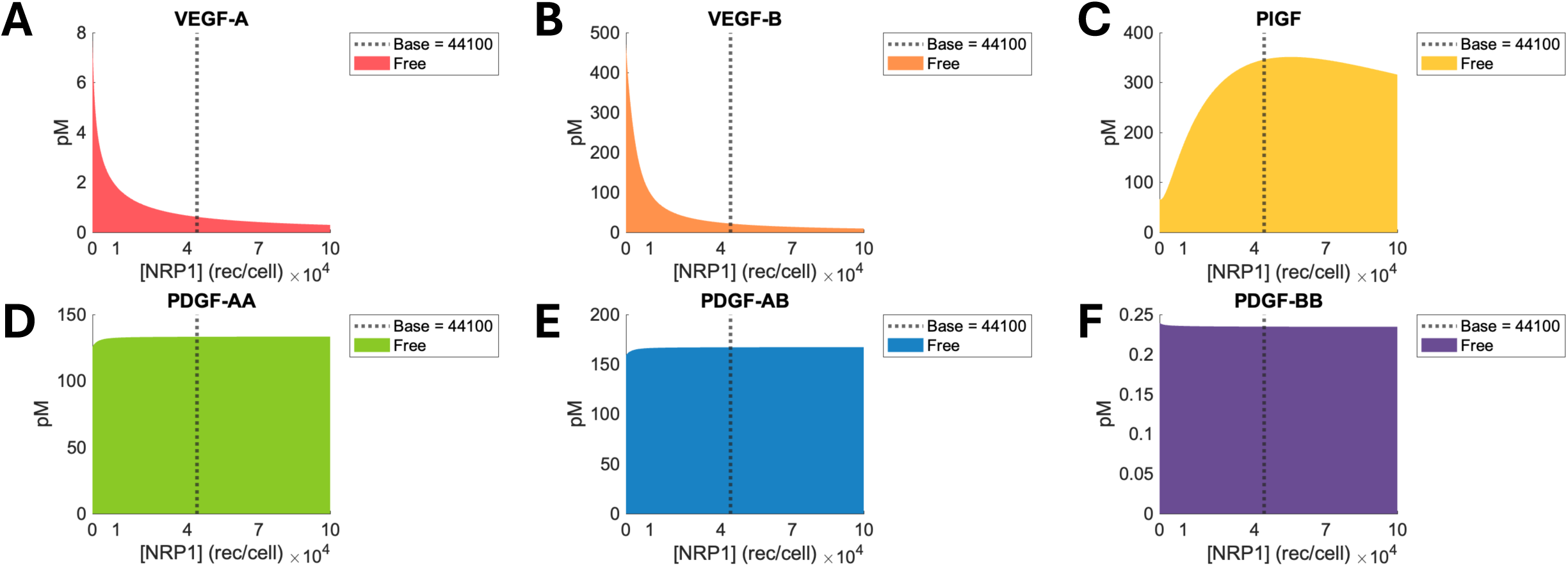
The concentration of free ligands at the equilibrium state depending on the various NRP1 densities. The colored area represents the concentration of free VEGF-A (A), VEGF-B (B), PlGF (C), PDGF-AA (D), PDGF-AB (E), and PDGF-BB (F) for each density of the receptor. The baseline of NRP1 density is 44,100 rec/cell.

**Fig 20.**
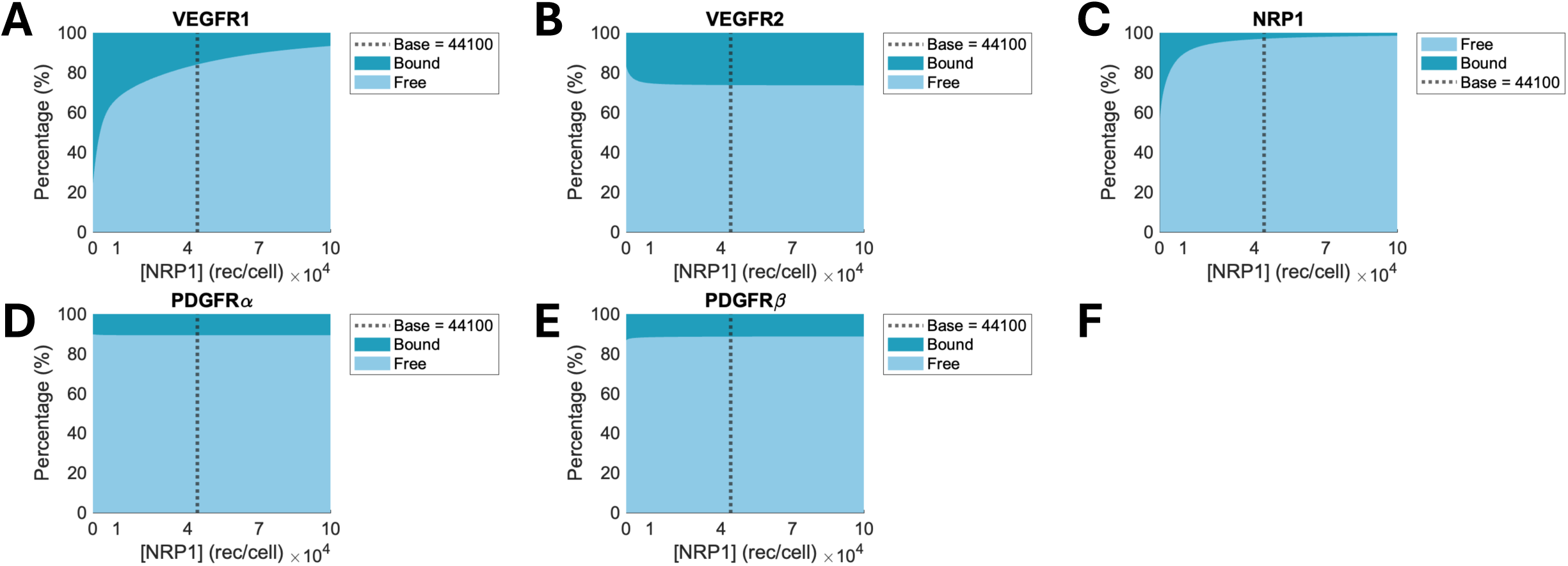
The percentage of free and bound receptors on the endothelial cell at the equilibrium state depending on the various NRP1 densities. The percentages of free and bound VEGFR1 (A), VEGFR2 (B), NRP1 (C), PDGFRα (D), and PDGFRβ (E) are represented. The light blue area represents the percentage of free receptors, while the dark blue area represents the percentage of bound receptors for each receptor density. The baseline of NRP1 density is 44,100 rec/cell.

**Fig 21.**
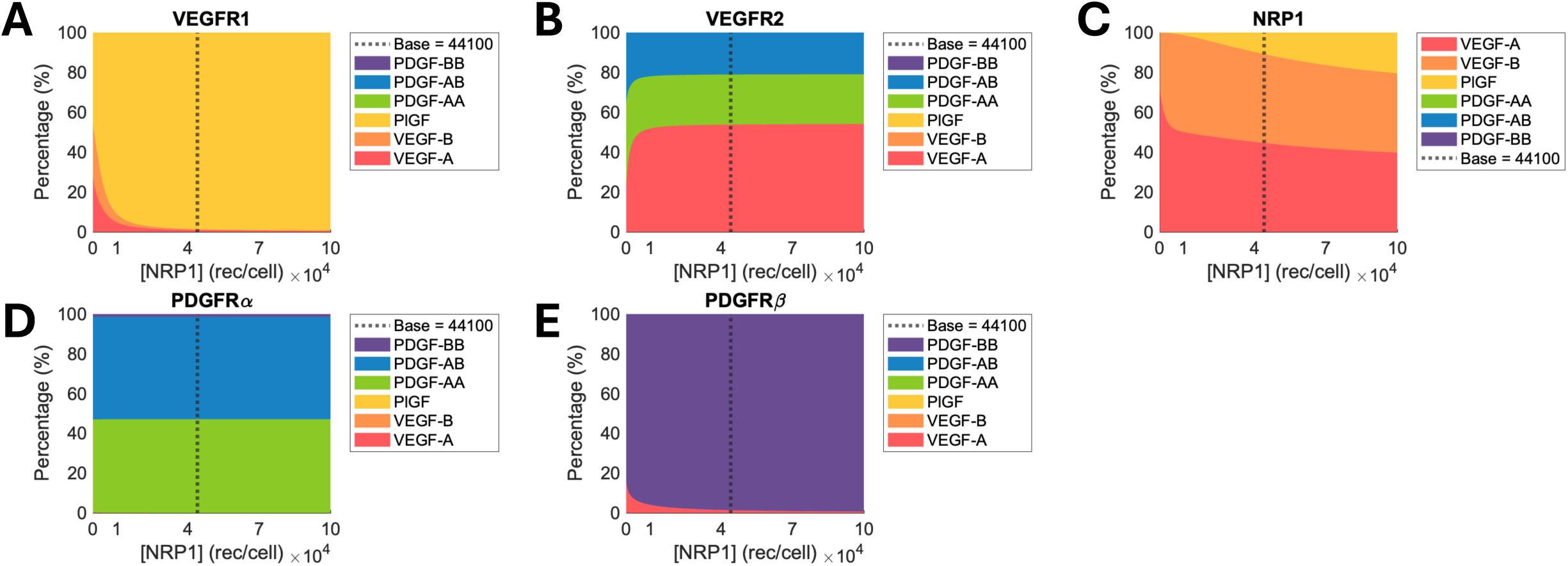
The percentage of ligands occupying each type of bound receptor at the equilibrium state depending on the various NRP1 densities. The percentages of VEGF-A, VEGF-B, PlGF, PDGF-AA, PDGF-AB, and PDGF-BB bound to VEGFR1 (A), VEGFR2 (B), NRP1 (C), PDGFRα (D), and PDGFRβ (E) are represented. The baseline value of the NRP1 density is 44,100 rec/cell.

**Fig 22.**
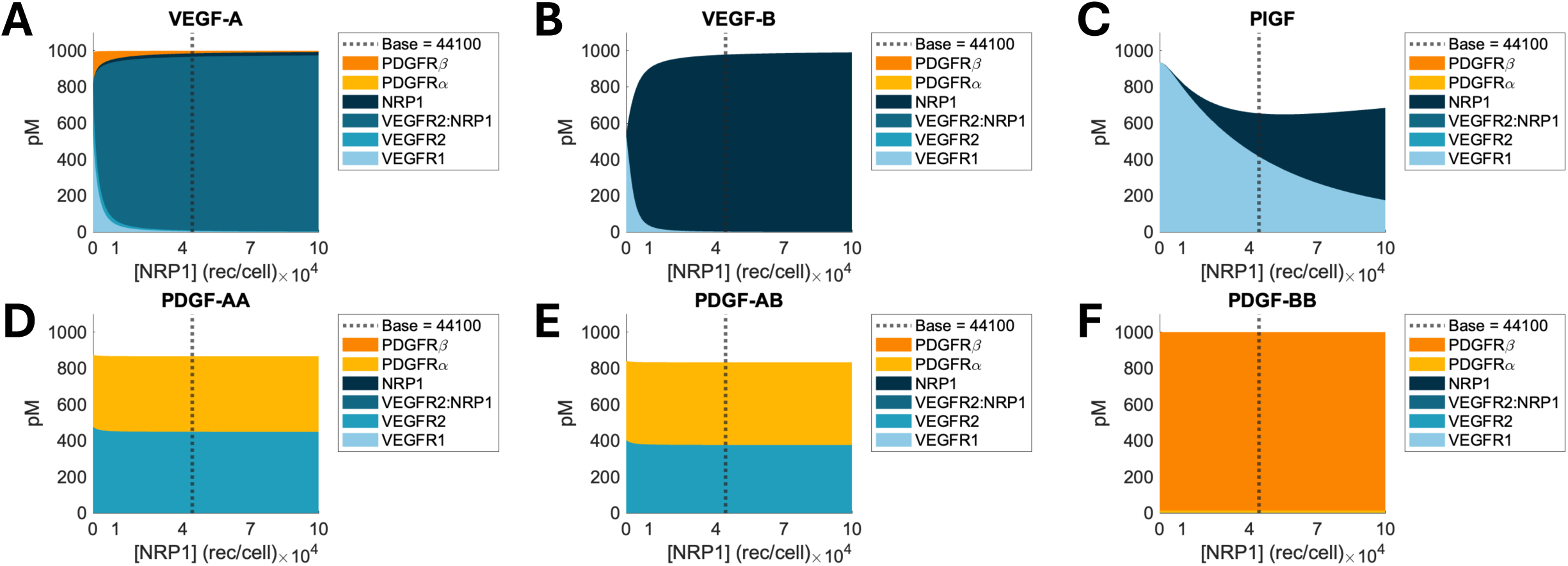
The concentration of ligand–receptor complex at the equilibrium state depending on the various NRP1 densities. The concentrations of VEGFR1, VEGFR2, NRP1, PDGFRα, and PDGFRβ that are bound to VEGF-A (A), VEGF-B (B), PlGF (C), PDGF-AA (D), PDGF-AB (E), and PDGF-BB (F) are represented. The baseline value of the NRP1 density is 44,100 rec/cell.

The increment in NRP1 density resulted in an increase in the formation of VEGF-A:NRP1, VEGF-A:VEGFR2:NRP1, and VEGF-B:NRP1 (Figs 22A–B). These increased formations led to five outcomes: 1) free VEGF-A and VEGF-B reduced by 7 pM and 450 pM, respectively (Figs 18A–B and 19A–B), 2) the percentage of unoccupied VEGFR2 reduced by 8% (Fig 20B), 3) the occupancy of PlGF in bound VEGFR1 increased by 49% (Fig 21A), 4) the occupancy of VEGF-A in bound VEGFR2 increased by 31% (Fig 21B), and 5) the occupancy of VEGF-A in PDGFRβ decreased by 14% (Fig 21E). The increased binding of VEGF family members to NRP1 led to an increase in unoccupied VEGFR1 by 70% (Fig 20A). The dynamics of free PDGFs and PDGFRs were not affected because PDGFR’s main binding partners, PDGFs, do not bind to NRP1 (Figs 18D–F, 19D–F, 20D–E, 21D, and 22D–F). Finally, the decreased percentage of bound NRP1 was due to the limited complex formation (Fig 20C).

### Increased PDGFRα density increases the formation of PDGFs:PDGFRα complexes

Our model confirmed that high PDGFRα density increased the PDGFs binding to PDGFRα. The increased PDGFs:PDGFRα formation resulted in four outcomes: 1) free VEGF-A, PDGF-AA, and PDGF-AB decreased (Figs 23A, 23D–E, 24A, and 24D–E), 2) the percentage of unoccupied VEGFR2 increased (Fig 25B) due to decreased PDGFs:VEGFR2 formation (Figs 27D–E), 3) the occupancy of VEGF-A in bound VEGFR2 increased (Fig 26B) because of maintained VEGF-A:VEGFR2:NRP1 or VEGF-A:NRP1 binding (Fig 27A) compared to reduced PDGFs:VEGFR2 binding (Figs 27D–E), and 4) the PDGF-BB:PDGFRα binding increased, while PDGF-BB:PDGFRβ binding decreased (Fig 27F). The increased PDGF-BB:PDGFRα binding increased the occupancy of PDGF-BB in PDGFRα (Fig 26D). The percentage of free PDGF-BB relatively unaffected compared to other PDGFs (Figs 23F and 24F) because the increased PDGF-BB:PDGFRα formation compensated for the decreased PDGF-BB:PDGFRβ formation (Fig 27F). The unoccupied PDGFRβ increased (Fig 25E) because of its decreased binding to PDGF-BB (Fig 27F). The ligand occupancy in PDGFRβ (Fig 26E) was unaffected because a majority of VEGF-A formed VEGF-A:VEGFR2:NRP1 or VEGF-A:NRP1 regardless of PDGFRα density (Fig 27A) and PDGF-BB has extremely stronger binding affinity than another binding partner, PDGF-AB.

**Fig 23.**
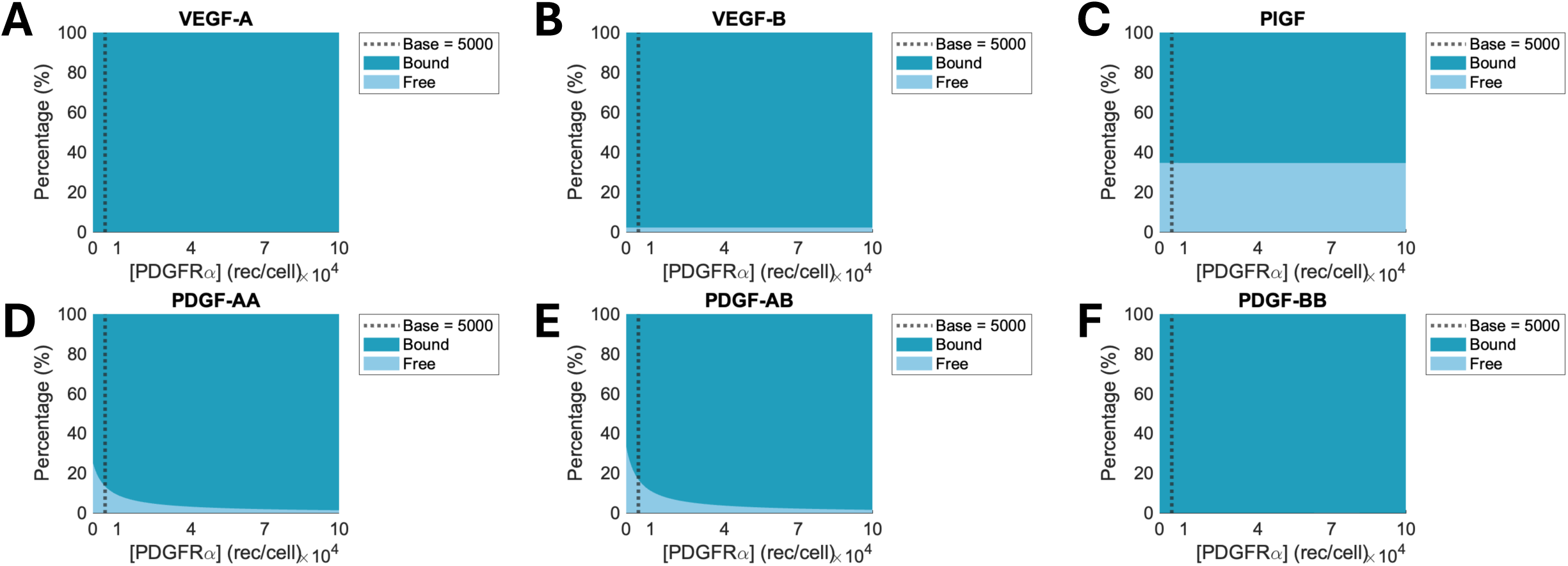
The percentage of free and bound ligands at the equilibrium state depending on the various PDGFRα densities. The percentages of the concentrations of free and bound VEGF-A (A), VEGF-B, (B), PlGF (C), PDGF-AA (D), PDGF-AB (E), and PDGF-BB (F) at each NRP1 density are plotted. The light blue area represents the percentage of free ligands, while the dark blue area represents the percentage of bound ligands for each receptor density. The baseline of PDGFRα density is 5,000 rec/cell.

**Fig 24.**
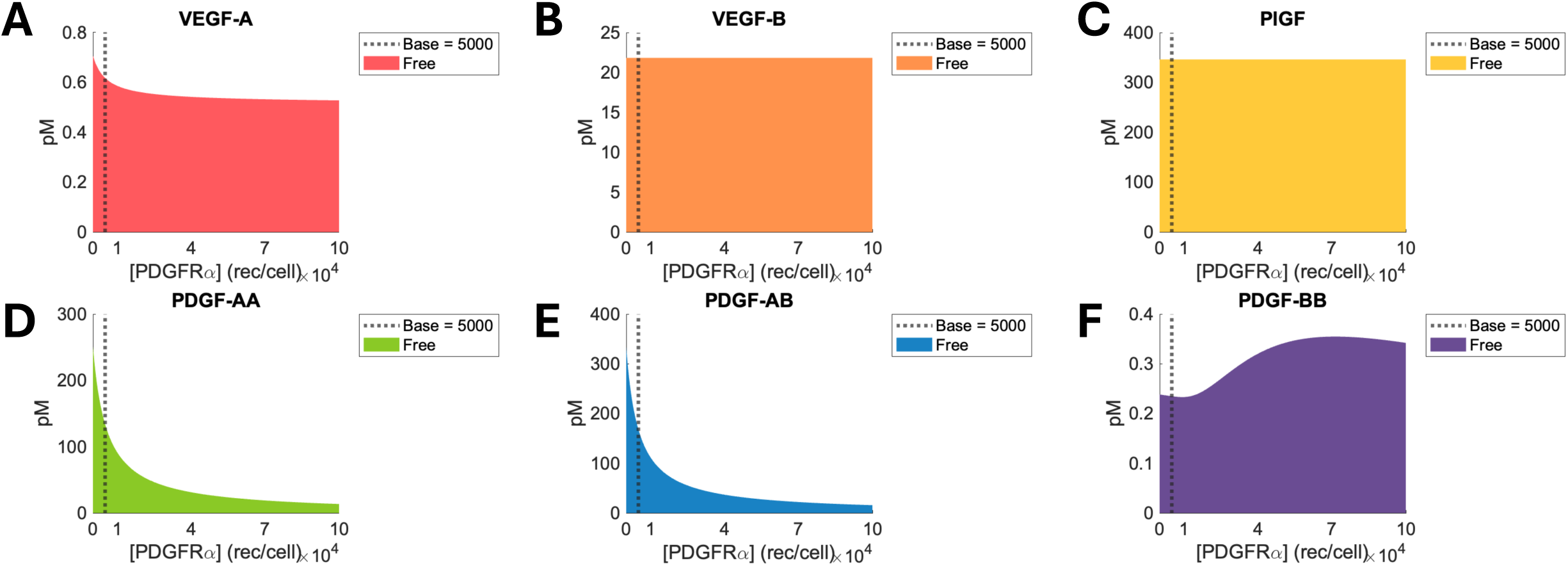
The concentration of free ligands at the equilibrium state depending on the various PDGFRα densities. The colored area represents the concentration of free VEGF-A (A), VEGF-B (B), PlGF (C), PDGF-AA (D), PDGF-AB (E), and PDGF-BB (F) for each density of the receptor. The baseline of PDGFRα density is 5,000 rec/cell.

**Fig 25.**
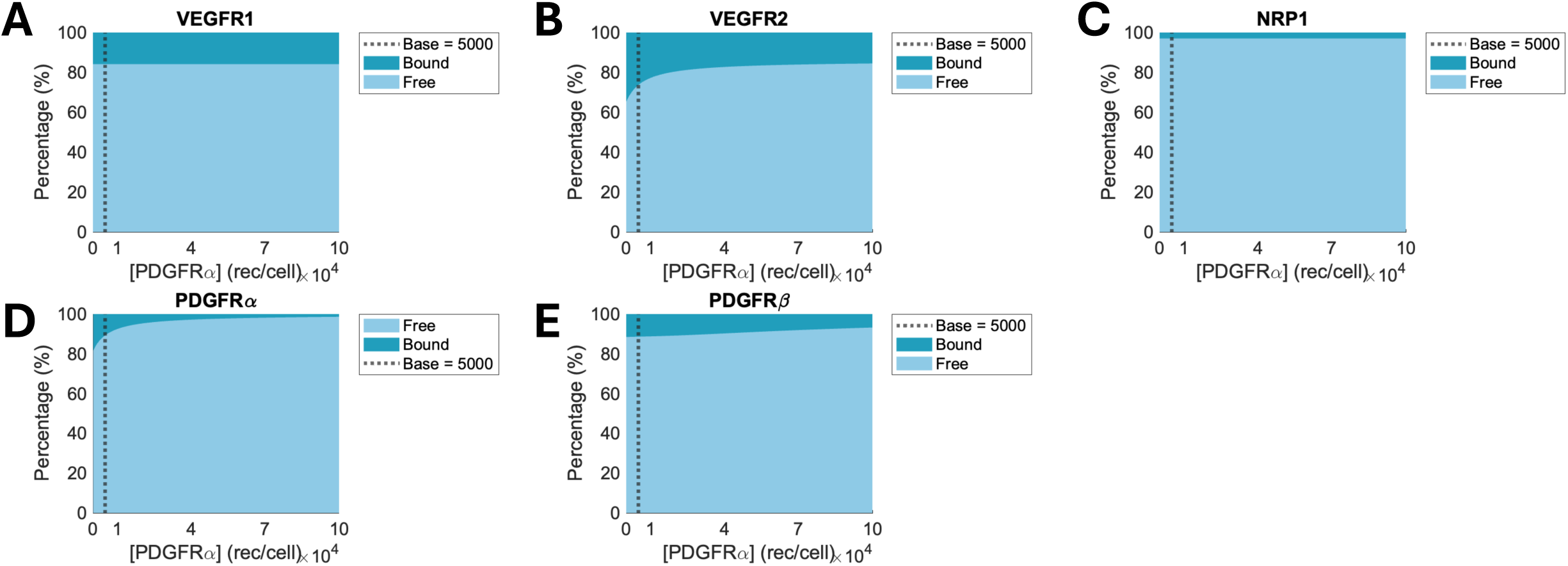
The percentage of free and bound receptors on the endothelial cell at the equilibrium state depending on the various PDGFRα densities. The percentages of free and bound VEGFR1 (A), VEGFR2 (B), NRP1 (C), PDGFRα (D), and PDGFRβ (E) are represented. The light blue area represents the percentage of free receptors, while the dark blue area represents the percentage of bound receptors for each receptor density. The baseline of PDGFRα density is 5,000 rec/cell.

**Fig 26.**
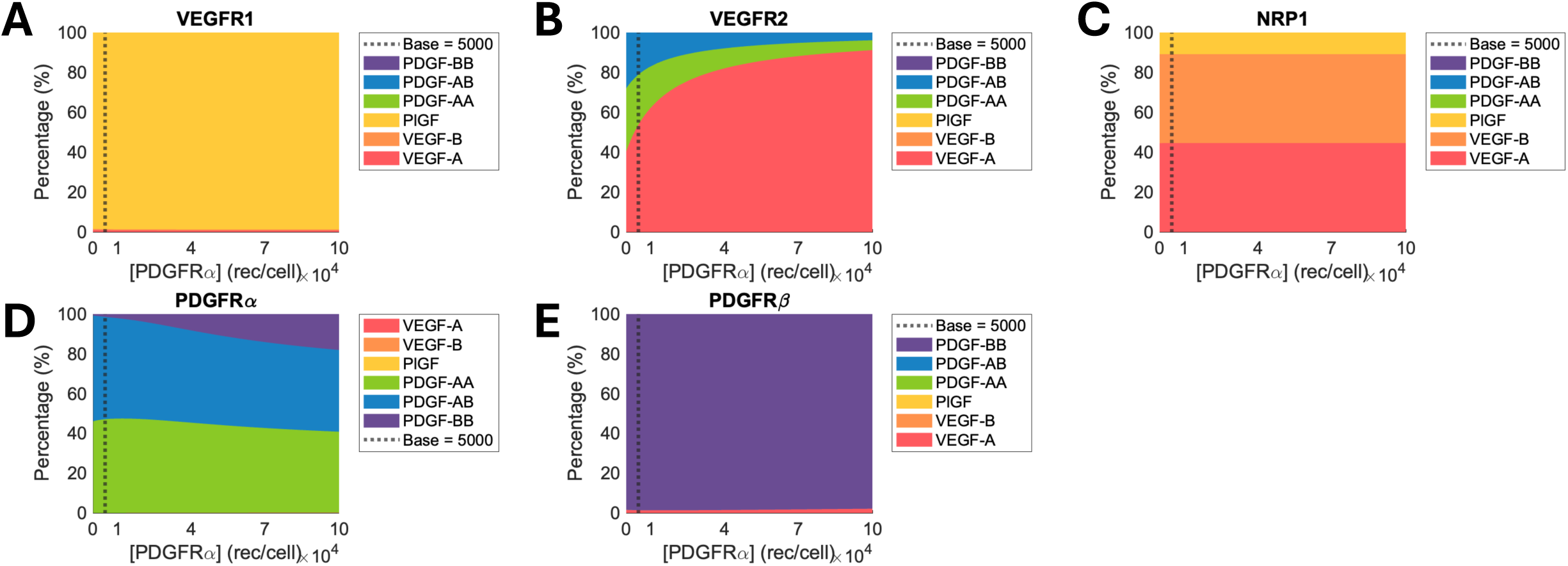
The percentage of ligands occupying each type of bound receptor at the equilibrium state depending on the various PDGFRα densities. The percentages of VEGF-A, VEGF-B, PlGF, PDGF-AA, PDGF-AB, and PDGF-BB bound to VEGFR1 (A), VEGFR2 (B), NRP1 (C), PDGFRα (D), and PDGFRβ (E) are represented. The baseline value of the PDGFRα density is 5,000 rec/cell.

**Fig 27.**
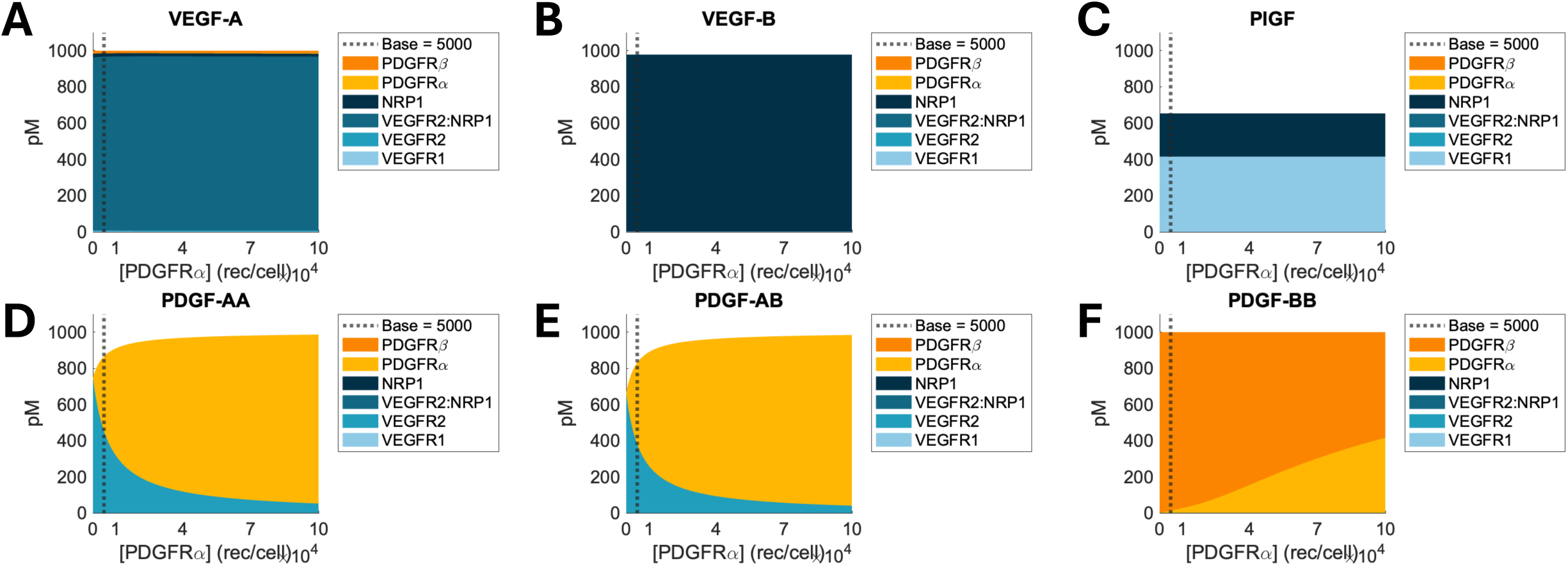
The concentration of ligand–receptor complex at the equilibrium state depending on the various PDGFRα densities. The concentrations of VEGFR1, VEGFR2, NRP1, PDGFRα, and PDGFRβ that are bound to VEGF-A (A), VEGF-B (B), PlGF (C), PDGF-AA (D), PDGF-AB (E), and PDGF-BB (F) are represented. The baseline value of the PDGFRα density is 5,000 rec/cell.

Increment in PDGFRα density affected neither the dynamics of unoccupied VEGFR1 and NRP1 (Figs 25A and 25C) nor ligand occupancy of bound VEGFR1 and NRP1 (Figs 26A and 26C). This was because their binding partners, VEGF-B and PlGF, do not bind to PDGFRα, thus, their dynamics were not affected by various PDGFRα density (Figs 23B–C, 24B–C, and 27B–C). Further, VEGF-A mainly formed VEGF-A:VEGFR2:NRP1 complexes (Fig 27A) as mentioned before. Finally, the increased unoccupied PDGFRα was observed as PDGFRα density increased, due to the limited complex formation (Fig 25D).

### Increased PDGFRβ density leads to increased VEGF-A:PDGFRβ

Our model predicted that the formation of VEGF-A:PDGFRβ increased and the formation of VEGF-A:VEGFR2:NRP1 decreased as PDGFRβ density increased (Fig 32A). This resulted in four additional outcomes: 1) free VEGF-A decreased by less than 0.2 pM (Figs 28A and 29A), 2) unoccupied VEGFR2 increased by 4% (Fig 30B), 3) the occupancy of VEGF-A in VEGFR2 and NRP1 decreased (Figs 31B–C), and 4) the occupancy of VEGF-A in PDGFRβ increased (Fig 31E). Increased formation of PDGF-BB:PDGFRβ complexes due to increased PDGFRβ density was accompanied by a decrease in free PDGF-BB (Figs 28F and 29F) and PDGF-BB:PDGFRα complex (Fig 32F). The reduction in PDGF-BB:PDGFRα led to an increase in unoccupied PDGFRα by 11% (Fig 30D) and a decrease in the occupancy of PDGF-BB in PDGFRα by 53% (Fig 31D) when PDGFRβ density increased from 0 rec/cell to 10^4^ rec/cell. The free PDGF-AA (Figs 28D and 29D) and PDGF-AA:VEGFR2 complex (Fig 32D) at the equilibrium state were not affected because PDGF-AA does not bind to PDGFRβ. The concentration of PDGF-AB:VEGFR2 and PDGF-AB:PDGFRα was also not affected (Fig 32E) while free PDGF-AB very slightly decreased by 4 pM due to PDGF-AB:PDGFRβ formation (Figs 28E and 29E). This was because PDGF-AB has an 800–1,000 times weaker binding affinity to PDGFRβ than VEGFR2 or PDGFRα.

**Fig 28.**
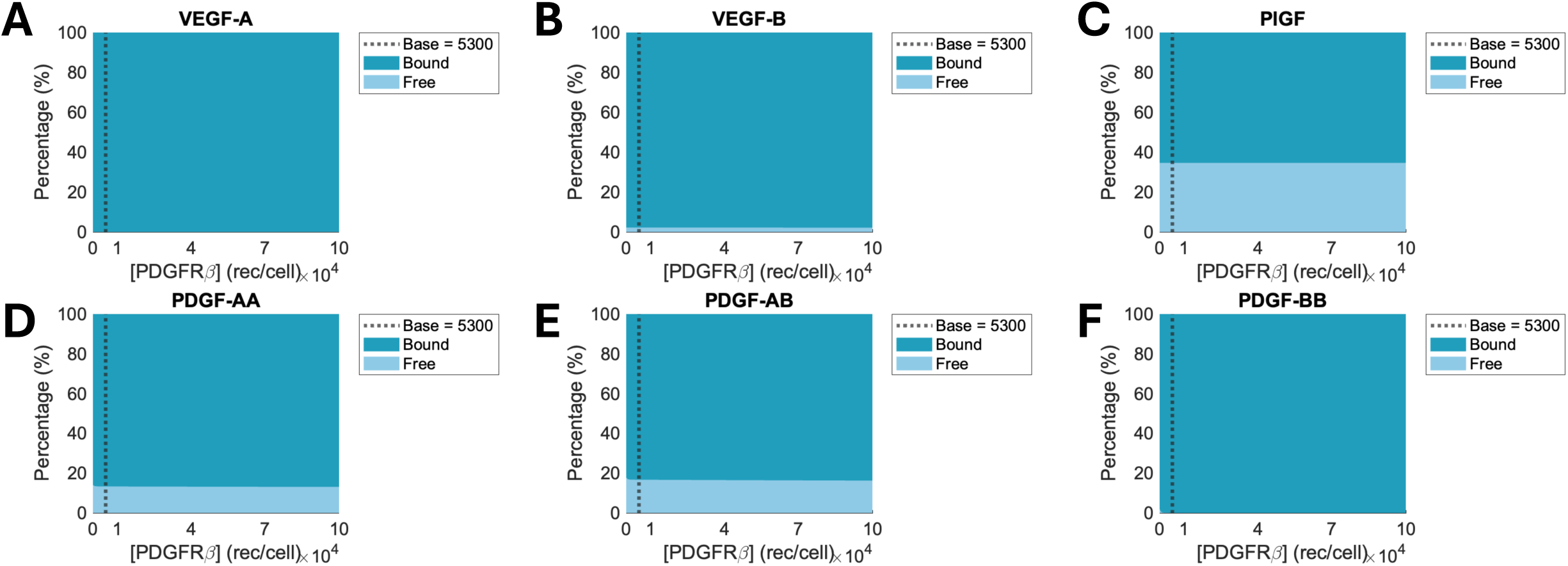
The percentage of free and bound ligands at the equilibrium state depending on the various PDGFRβ densities. The percentages of the concentrations of free and bound VEGF-A (A), VEGF-B, (B), PlGF (C), PDGF-AA (D), PDGF-AB (E), and PDGF-BB (F) at each NRP1 density are plotted. The light blue area represents the percentage of free ligands, while the dark blue area represents the percentage of bound ligands for each receptor density. The baseline of PDGFRβ density is 5,300 rec/cell.

**Fig 29.**
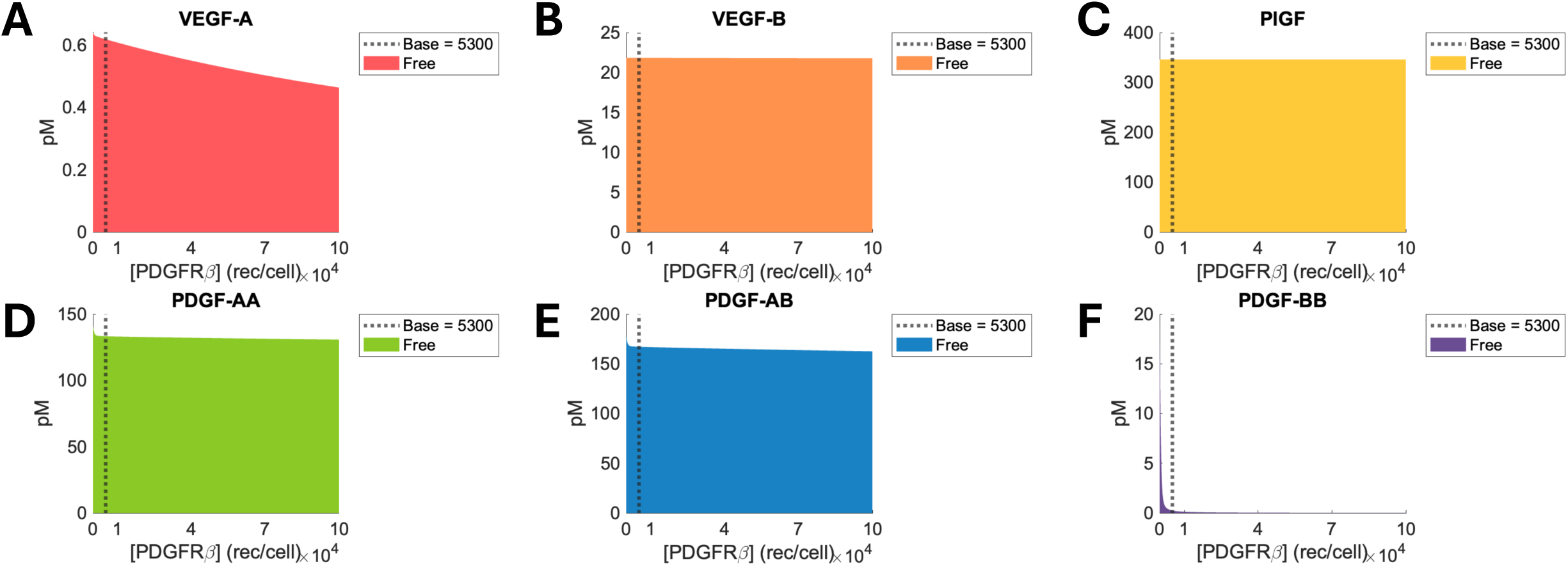
The concentration of free ligands at the equilibrium state depending on the various PDGFRβ densities. The colored area represents the concentration of free VEGF-A (A), VEGF-B (B), PlGF (C), PDGF-AA (D), PDGF-AB (E), and PDGF-BB (F) for each density of the receptor. The baseline of PDGFRβ density is 5,300 rec/cell.

**Fig 30.**
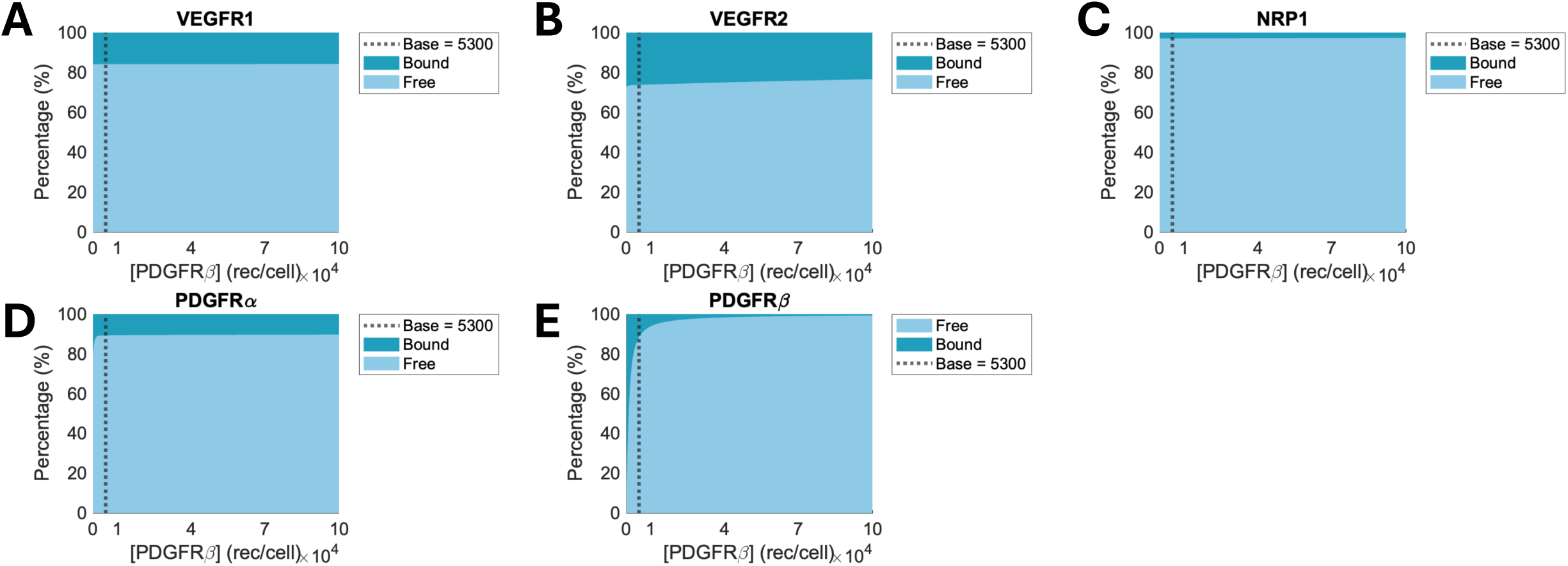
The percentage of free and bound receptors on the endothelial cell at the equilibrium state depending on the various PDGFRβ densities. The percentages of free and bound VEGFR1 (A), VEGFR2 (B), NRP1 (C), PDGFRα (D), and PDGFRβ (E) are represented. The light blue area represents the percentage of free receptors, while the dark blue area represents the percentage of bound receptors for each receptor density. The baseline of PDGFRβ density is 5,300 rec/cell.

**Fig 31.**
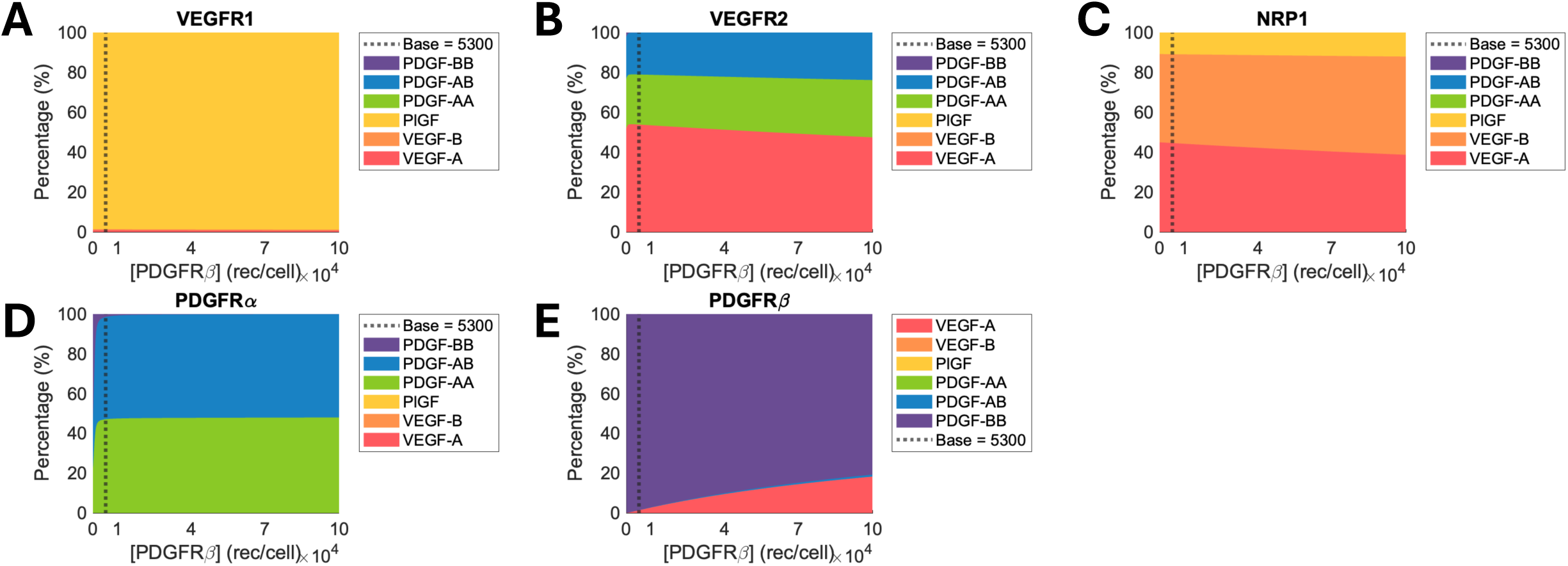
The percentage of ligands occupying each type of bound receptor at the equilibrium state depending on the various PDGFRβ densities. The percentages of VEGF-A, VEGF-B, PlGF, PDGF-AA, PDGF-AB, and PDGF-BB bound to VEGFR1 (A), VEGFR2 (B), NRP1 (C), PDGFRα (D), and PDGFRβ (E) are represented. The baseline value of the PDGFRβ density is 5,300 rec/cell.

**Fig 32.**
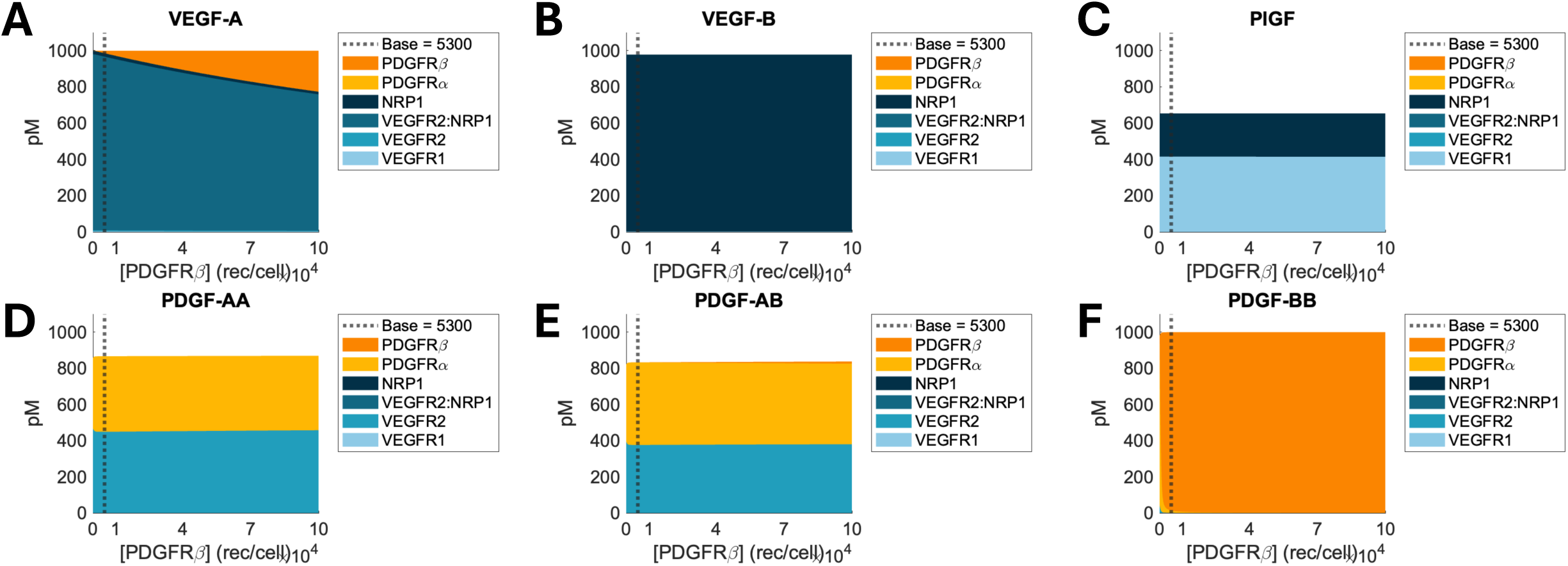
The concentration of ligand–receptor complex at the equilibrium state depending on the various PDGFRβ densities. The concentrations of VEGFR1, VEGFR2, NRP1, PDGFRα, and PDGFRβ that are bound to VEGF-A (A), VEGF-B (B), PlGF (C), PDGF-AA (D), PDGF-AB (E), and PDGF-BB (F) are represented. The baseline value of the PDGFRβ density is 5,300 rec/cell.

Our model predicted that unoccupied VEGFR1 and NRP1 and the ligand occupancy in VEGFR1 were not affected (Figs 30A, 30C, and 31A) for three reasons: 1) their binding partners, VEGF-B and PlGF, do not bind to PDGFRβ, thus, maintaining their dynamics at the equilibrium state (Figs 28B–C, 29B–C, and 32B–C), 2) the concentration of VEGF-A:VEGFR1 complex was very low regardless of PDGFRβ density (Fig 32A), and 3) the unoccupied NRP1 increased due to 200 pM of decreased concentration of VEGF-A:NRP1 and VEGF-A:VEGFR2:NRP1 (Fig 32A), which was much smaller than NRP1 density (73 nM). Finally, the increase in the percentage of unoccupied PDGFRβ was due to the limited complex formation compared to the increased amount of PDGFRβ density (Fig 30E).

## Discussion

In this study, we established a computational model describing cross-family interactions on an endothelial cell surface based on mass-action kinetics. The computational model considers three VEGFs (VEGF-A, VEGF-B, and PlGF) and three PDGF family members (PDGF-AA, PDGF-AB, and PDGF-BB). Here, we identified which ligand dominantly binds to which receptors, since both families compete to bind overlapping binding partners and the signaling properties of each complex are different. Our model offers quantitative evidence about how different binding kinetics of VEGF and PDGF families regulate the formation of complexes under the same ligand concentration.

### Comparisons of kinetics data in our study with previous PDGF binding studies

We conducted the SPR assay to measure lacked association and dissociation kinetic data. Our study showed that PlGF binds to VEGFR1 with a strong binding affinity of 46.5 pM. This was five-times stronger than reported binding affinity of radiolabeled ^125^I-PlGF to VEGFR1 on HUVEC as 230 pM in previous studies [69]. Our SPR assay also showed that VEGF-B binds to VEGFR1 with the binding affinity of 579 pM, which was also five times larger than the previous measurements, 114 pM [70].

Our study found that PDGF binds to VEGFR2 with binding affinity of 1–2 nM. Although these kinetic data was previously established by Mamer et al. [41], we reexamined these data for two reasons: 1) the previously determined data were from old SPR machine, and 2) reestablishing kinetics data will allow us to check if our measurements align with previous data. Indeed, Mamer et al. showed a very large range of PDGF:VEGFR2 binding ranging from 110 pM to 530 nM (530 nM for PDGF-AA:VEGFR2, 110 pM for PDGF-AB:VEGFR2, and 37 nM for PDGF-BB:VEGFR2). Furthermore, while our study showed 2 nM of binding affinities for PDGF:PDGFRs except for PDGF-BB:PDGFRs, which was in the picomolar range, the values reported by Mamer et al. were in the range from 17 nM to 2.2 µM. These large variations may be attributed to the low sensitivity of the old machine. We could not measure the binding affinity of PDGF-AB:PDGFRβ, which may be attributed to PDGF-AB very low affinity to PDGFRβ (K_d_ = 2.23 µM [41]. Overall, we chose newly measured kinetics data as they were more similar to other canonical interactions like VEGF-A:VEGFR2 (K_d_ = 167 pM) than previously established values.

### Role of PlGF as the main binding partner of VEGFR1 in computational models and pathological conditions

Setting a uniform ligand concentration allowed us to examine the differential binding dynamics, here PlGF emerged as the dominant binding partner of VEGFR1. It occupied over 95% of bound VEGFR1 despite the weaker binding affinity than VEGF-A:VEGFR1. This outcome resulted from three factors determining the formation of VEGF-A complexes: 1) the concentration of free VEGF-A, 2) association rate constants of VEGF-A to VEGFR1, VEGFR2, and NRP1, and 3) densities of the receptors. Although VEGF-A:VEGFR1 has 2.4– and 8.5-times higher association rate constants than VEGFR2 and NRP1, respectively, the densities of VEGFR2 and NRP1 are 3 and 28 times higher than VEGFR1. Finally, the receptors share the same VEGF-A concentration. Thus, the relatively low association rate of VEGFR2 and NRP1 is compensated for by their high densities during complex formation. Besides, the coupling of VEGFR2 and NRP1 stabilizes the binding of VEGF-A to these receptors. Therefore, VEGF-A predominantly binds to VEGFR2 and NRP1 rather than VEGFR1. VEGF-B preferentially binds to NRP1 because of a similar reason. These results were consistent with previous studies. Clegg et al. demonstrated that VEGF-A165 was the main binding partner of VEGFR2 in their whole-body model focusing on the calf muscle [57]; VEGF-A165:VEGFR2 complex occupied 72% of total VEGFR2 ligation. On the other hand, PlGF:VEGFR1 occupied 29% of VEGFR1 ligation, following 70% occupation of VEGF-A121:VEGFR1:NRP1. VEGF-A165:VEGFR1 occupied only 1%. Since no competitive binding was observed in the model, if VEGF-A121 were removed, it is expected that PlGF would occupy the most VEGFR1 ligation, as shown in our model.

To further support our finding on PlGF as a dominant regulator of VEGFR1 signaling, we set the PlGF concentration as 0 nM and performed the simulation again (Fig S3). Our computational model showed that in the absence of PlGF, VEGFR1 primarily binds to VEGF-A and VEGF-B instead of PlGF (Fig S3A). Interestingly, the absence of PlGF does not appear to impact VEGF-A binding to VEGFR2, and thus VEGFR2 signaling likely remains unaffected. Specifically, the percentages of ligands bound to VEGFR2 were conserved even in the absence of PlGF (Fig 3 vs. Fig S3A). Furthermore, the percentage of bound VEGFR2 was not affected while the percentage of bound VEGFR1 decreased (Fig S3B). This model prediction is consistent with other experimental data, where PlGF-knockdown human retinal endothelial cells showed markedly reduced VEGFR1 phosphorylation while VEGFR2 phosphorylation remained unchanged [71]. PlGF is usually expressed in the placenta, but cancer cells or retinal pigment epithelial cells in diabetic patients also overexpress PlGF under hypoxic conditions [72–74]. These results suggest that VEGFR1 signaling is mainly regulated by PlGF when it is present, including in pathological conditions.

**Fig S3.**
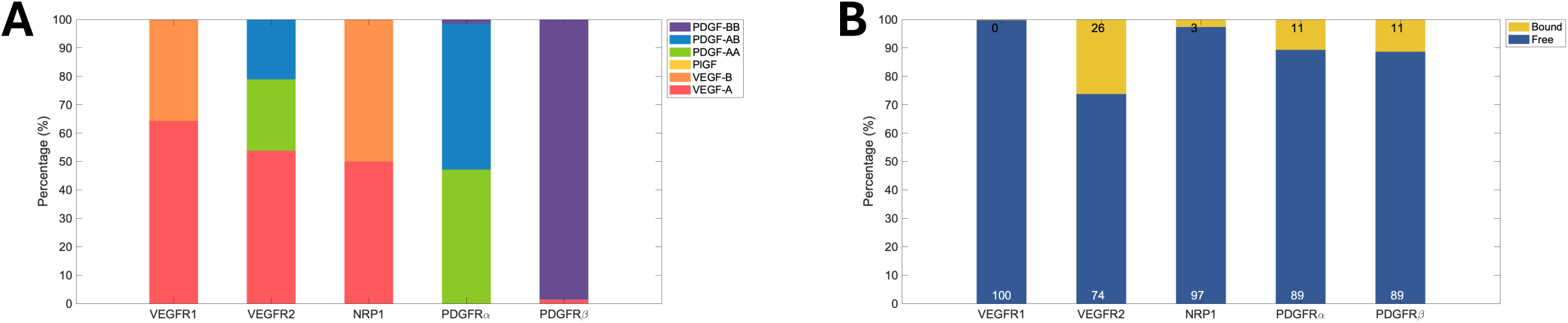
Changes in cross-family interactions on endothelial cells when PlGF is removed. (A) The percentages of VEGF-A, VEGF-B, PlGF, PDGF-AA, PDGF-AB, and PDGF-AB that occupy each bound receptor are shown in red, orange, yellow, green, blue, and purple, respectively. (B) The percentages of free and bound receptors are presented.

### Finding that PDGFs occupy only 50% of VEGFR2 is inconsistent with previous computational models

Our result showed that 50% of VEGFR2 is occupied by PDGF-AA and PDGF-AB when the ligand concentrations are 1 nM, which was not consistent with the previous studies. Mamer *et al.* (2017) predicted that PDGF isoforms occupy 96% of VEGFR2 ligation from a computational model for interactions of VEGF-A and PDGF isoforms (PDGF-AA, PDGF-AB, and PDGF-BB) with VEGFR2 [41]. This difference may be attributed to the different PDGF concentrations used in the study. Indeed, Mamer et al. used the ligand concentration in the serum of healthy humans, and the concentration of VEGF-A was 3–100 times lower than PDGF isoforms (VEGF-A: 81.7–91.83 pg/ml; PDGF-AA: 250.0–1,700 pg/ml; PDGF-AB: 250.0–2,830 pg/ml; PDGF-BB: 8,506±550 pg/ml). Even though the binding affinity of VEGF-A:VEGFR2 is 9–12 times stronger than those of PDGFs:VEGFR2, the higher concentration of PDGFs would elevate the formation of PDGFs:VEGFR2. Thus, if we apply experimentally measured concentration of ligands to our model, it is expected that the percentage of PDGF isoforms occupying VEGFR2 would increase.

### VEGFR1 overexpression resulting in reduced VEGFR2 complex does not mean low angiogenic signal

Our model showed that an increase in VEGFR1 density increased the binding of ligands to VEGFR1 while it decreased the concentration of VEGF-A:VEGFR2. Also, the percentage of PDGFs occupying VEGFR2 increased due to increased interactions between PDGFs and VEGFR2. One might think that overall angiogenic activity would be lowered with increasing VEGFR1 density because of the following two reasons: 1) VEGFR1 binding to its ligands shows weaker signaling on endothelial cells than VEGF-A:VEGFR2 complex [14], and 2) PDGF:VEGFR2 cannot exhibit angiogenic signal because PDGFs cannot phosphorylate VEGFR2 (Liu et al., unpublished data). However, increasing evidence suggests that binding ligands to the high density of VEGFR1 may compensate for the low efficiency of VEGFR1 signaling and enhance the cell response. For example, the overexpression of VEGFR1 on endothelial cells has been reported in several pathological conditions, including tumors, obesity, and hindlimb ischemia [51,75,76], which requires angiogenesis. The VEGFR1 density on the endothelial cell surface in tumors has been reported to be 7–8 times larger than in normal tissues [44,52,75]. Also, the computational models for VEGFR1–overexpressing tumor have predicted the reduced effect of anti-VEGF on VEGF level in plasma [44]. Furthermore, when PlGF-overexpressing tumor cells were inoculated in mice lacking VEGFR1 tyrosine domain, the tumor growth and vessel formation were significantly slower than in the wild-type mice. Our result showing the higher occupation of PlGF than VEGF-A and VEGF-B for VEGFR1 density in the range of VEGFR1 density reported in pathological conditions may imply that PlGF:VEGFR1 signaling plays a significant role in pathological angiogenesis, supporting this experimental data.

### Increase in VEGF-A:VEGFR2:NRP1 complex by VEGFR2 overexpression is inconsistent with previous computational studies for peripheral arterial disease, and increase in PDGF:VEGFR2 may related to low angiogenesis

Our model showed increased VEGF-A:VEGFR2:NRP1 formation for increased VEGFR2 density. This result was inconsistent with previous computational studies. For example, the impact of overexpression of VEGFR2 on complex formation was investigated in computational studies for peripheral arterial disease (PAD) [77]. In a computational model developed by Wu et al., the study showed that an increase in VEGFR2 density resulted in the formation of VEGF-A:VEGFR2 complex becoming dominant over VEGF-A:VEGFR2:NRP1 complex formation. This difference was due to the different ratios of VEGFR2 and NRP1. In our model, the ratio of VEGFR2 and NRP1 densities was 1:9, while it was 1:0.2 in the PAD model. The higher NRP1 level compared to VEGFR2 in our model made a big pool of unoccupied NRP1 available regardless of the VEGFR2 density changes.

In patients suffering from PAD, increased VEGFR2 expression has been reported despite low level of angiogenesis in the diseased tissue [78]. Furthermore, both our model and Wu et al.’s model predicted a high concentration of VEGFR2 complex bound to VEGF-A. Wu et al. suggested two possibilities: 1) VEGFR2 downstream signaling is interrupted, and 2) soluble VEGFR1 inhibits VEGFR2 signaling by dimerizing with VEGFR1 or VEGFR2 monomer. However, our model suggests that PDGFRα may play a role in PAD. Indeed, the bindings of PDGF-AA and PDGF-AB to PDGFRα decreased while their binding to VEGFR2 increased. The increase in VEGF-A:VEGFR2 binding was not noticeable because most VEGF-A was bound to VEGFR2 even at the low VEGFR2 density level. Considering that PDGF cannot phosphorylate VEGFR2, the reduced percentage of bound PDGFRα may result in reduced angiogenesis in the muscles in PAD patients.

### Necessity of developing tissue or whole-body level computational models because of inconsistent results that free PlGF decreases upon NRP1 overexpression

Our study showed that the concentration of free PlGF was increased when the NRP1 density was below the baseline, and then it slightly decreased as NRP1 density was further increased. Considering that the baseline value of NRP1 density is from normal endothelial cells, our result was inconsistent with previous experimental studies because the high NRP1 density in several cancer cell lines [9,79] and the higher PlGF level in plasma in cancer patients than healthy subjects have been reported [80,81]. Further, it has been shown that both PlGF mRNA and protein expressions are upregulated in tumors [82,83]. Therefore, developing computational models for cross-family interactions at the tissue or whole-body level is required to investigate the relation between free PlGF concentration and NRP1 overexpression.

### Increased PDGF-BB:PDGFRα due to PDGFRα overexpression may be related to tumor growth

Our study showed that increased PDGFRα density reduced PDGFs:VEGFR2 complexes and increased the formation of the PDGFs:PDGFRα complexes. Specifically, high PDGFRα density increased the PDGF-BB binding to PDGFRα. This result may imply enhanced cell signaling because 1) it was reported that PDGF bound VEGFR2 cannot generate cell signaling, as mentioned above (Liu et al., unpublished data), and 2) PDGF-BB is considered to be more potent than PDGF-AA or PDGF-AB in PDGFRα activation (7-fold higher PDGF-BB induced PDGFRα phosphorylation compared to PDGF-AA) [84,85]. Some evidence also supports the enhanced cell signaling due to increased PDGFRα density: 1) PDGFRα was shown to be overexpressed in some types of tumors and associated with metastasis [86–88], and 2) PDGF-BB-induced PDGFRα activation contributes to the tumor invasion via MAPK/ERK and PI3K/Akt pathways [88], which are also known to regulate cell proliferation and cell survival [89,90]. Our computational model might be able to provide an explanation of why PDGFRα-overexpressing tumors exhibit invasiveness.

### Increased VEGF-A:PDGFRβ due to overexpressed PDGFRβ may be related to altered angiogenesis in tumor and adipose tissue

Our model predicted that overexpression of PDGFRβ on endothelial cell surface would increase the VEGF-A:PDGFRβ formation while it would decrease VEGF-A:VEGFR2 formation. Although there is a lack of information to decide if VEGF-A:VEGFR2 exhibits a stronger signal than VEGF-A:PDGFRβ, our result may indicate the altered angiogenesis based on the following evidence: 1) the PDGFRβ-overexpressing tumors were not necessarily correlated with metastasis [88], and 2) PDGFRβ expression was upregulated in aged adipose tissue, and PDGFRβ signaling suppressed the beige adipose tissue development [91]. Although the study suggested that the reduced beige adipogenesis may be due to the activation of cold-induced immune cells, an increase in VEGF-A:PDGFRβ formation might also be attributed to the failure of beige adipogenesis.

### Our effort to ensure the reproducibility of our computational model

The reproducibility of computational models in systems biology is an important factor for advances in science. However, many computational studies do not fully describe the simulation settings or how the model was developed, interfering with reproducibility. Three requirements for reproducible modeling were raised by Medley et al. (2016) [92]: 1) researchers should clarify the data source and model assumptions, 2) researchers should clarify parameter values, the algorithm, and options of the simulation software they used, and 3) researchers should check if statistical outcomes from several software tools are the same. Our study successfully reproduced previously well-established computational model and validated our model by comparing outcomes of both models. Furthermore, our study provided the references of each parameter value and initial concentration of species, and the model assumptions were represented in the Method sections. We described the parameter values, the integration algorithm used to solve the system of differential equations, the tolerance for local errors in the integration, and the software version to ensure reproducibility. Furthermore, we transcribed a system of ordinary differential equations in Supplementary material. Finally, since our model is deterministic, not stochastic, there are fewer concerns about differences in statistical outcomes. Overall, we believe that our computational model is reproducible.

### Future studies

We developed a computational model including the binding of ligands and receptors and receptor couplings but did not include the secretion or degradation rate of ligands or the trafficking process of receptors. Considering that the equilibrium state in our model was reached ten days after the simulation start time, the secretion and degradation of ligands and receptors would influence the dynamics of cross-family interactions.

Also, including the internalization rate of receptors would change the time-dependent profile of complex formations since the ligand having low binding affinity may not have enough time to bind its receptors. Thus, only ligands showing strong binding affinities may mediate angiogenesis as illustrated in Fig 4, where we recapitulated the PDE model established by Mac Gabhann and Popel. Thus, although we excluded those trafficking parameters in this study to focus on the influence of cross-family interactions on complex formation, our future work will include those trafficking processes.

Although our model did not include heterodimers due to the lack of information about how they are involved in cross-family interactions. Heterodimers, especially between VEGFRs and PDGFRs, may provide another possibility of cross-family interaction. An example of this possibility has already been found: heme oxygenase-1 promotes the formation of inactive heterodimers of VEGFR2 and PDGFRβ in vascular smooth muscle cells, diminishing PDGFRβ signaling [93]. With the aid of molecular dynamics simulations and AI tools like AlphaFold, future studies would identify additional, energetically favorable heterodimeric receptor complexes. Incorporation of this knowledge into future biological experiments and computational modeling would expand our understanding of angiogenesis.

Our model did not consider VEGF-C and VEGF-D, which are lymphangiogenesis regulators. These growth factors bind to VEGFR2 and VEGFR3, thus, including them in the model would affect model outcomes such as the number of VEGFR2 complexes. Similarly, PDGF-CC and PDGF-DD bind to PDGFRα and PDGFRβ, respectively. Including these PDGFs would alter the dynamics of PDGFRs and their ligand distribution at the equilibrium state. Therefore, future work should include these growth factors.

Finally, our model assumed the identical concentration (1 nM) for all ligands, but the actual serum ligand levels are different in the human body. As examined by Mamer *et al.*, the concentration of VEGF-A is 3–100 times lower than PDGF isoforms (82 pg/ml for VEGF-A and 250 pg/ml or 8,506 pg/ml for PDGF-AA, PDGF-AB, and PDGF-BB) [41], and PlGF plasma level and VEGF-B serum level were reported around 12.5 pg/ml and 116 pg/ml in human, respectively [94,95]. The implication of different ligand concentrations would affect the overall dynamics illustrated in this paper. Thus, our future work will include the application of physiological ligand concentration.

## Conclusion

We developed a computational model to investigate cross-family interactions between VEGF and PDGF families. The model parameters were derived through a combination of literature search and surface plasmon resonance assays. Our model was validated by comparing outcomes with a previously well-established model under the same parametrical settings [61]. Furthermore, our model predicts the distribution of ligands bound to each receptor under the same ligand concentrations.

Our study contributes to advancing our understanding in three key areas: 1) **Computational modeling**: our model establishes a fundamental framework for exploring cross-family signaling on endothelial cell surfaces. This framework can be extended to model signaling processes at any tissue level or within intracellular molecule networks. 2) **Bioengineering field**: our study provides new insight into which ligand predominantly occupies each receptor in angiogenesis. Understanding this mechanism is important as different ligand-receptor complexes play distinct roles in angiogenesis. 3) **Vascular field**: the study offers potential avenues for developing therapeutic strategies targeting multiple growth factors to treat pathological conditions. Understanding ligand distribution across receptors can elucidate the relative contributions of each ligand to receptor-mediated signaling and help determine appropriate treatment targets.

## Acknowledgments

Author contributions

PII, YL, and YF formulated the research goals and aims. YL developed the computational model and implemented computational algorithms. SK performed experiments and analyzed the data. All authors wrote the original draft preparation.

## Data availability statement

The data and code supporting the findings of this study are available within this paper, its supplementary file, and at: https://github.com/YunjeongLee/x-family_membrane.git.

