## Supplementary File for "Cross-family interactions of vascular endothelial growth factors and platelet-derived growth factors on the endothelial cell surface: A computational model"

**Glossary**

| **Symbol** | **Interpretation** |
| --- | --- |
| [VA] | The concentration of VEGF-A |
| [VB] | The concentration of VEGF-B |
| [Pl] | The concentration of PlGF |
| [PDAA] | The concentration of PDGF-AA |
| [PDAB] | The concentration of PDGF-AA |
| [PDBB] | The concentration of PDGF-BB |
| [R1] | The concentration of unoccupied VEGFR1 |
| [R2] | The concentration of unoccupied VEGFR2 |
| [N1] | The concentration of unoccupied NRP1 |
| [PDRa] | The concentration of unoccupied PDGFRα |
| [PDRb] | The concentration of unoccupied PDGFRβ |
| [VA:R1] | The concentration of VEGF-A:VEGFR1 complex |
| [VA:R2] | The concentration of VEGF-A:VEGFR2 complex |
| [VA:R2:N1] | The concentration of VEGF-A:VEGFR2:NRP1 complex |
| [VA:N1] | The concentration of VEGF-A:NRP1 complex |
| [VA:PDRa] | The concentration of VEGF-A:PDGFRα complex |
| [VA:PDRb] | The concentration of VEGF-A:PDGFRβ complex |
| [VB:R1] | The concentration of VEGF-B:VEGFR1 complex |
| [VB:N1] | The concentration of VEGF-B:NRP1 complex |
| [Pl:R1] | The concentration of PlGF:VEGFR1 complex |
| [Pl:N1] | The concentration of PlGF:NRP1 complex |
| [PDAA:R2] | The concentration of PDGF-AA:VEGFR2 complex |
| [PDAA:PDRa] | The concentration of PDGF-AA:PDGFRα complex |
| [PDAB:R2] | The concentration of PDGF-AB:VEGFR2 complex |
| [PDAB:PDRa] | The concentration of PDGF-AB:PDGFRα complex |
| [PDAB:PDRb] | The concentration of PDGF-AB:PDGFRβ complex |
| [PDBB:R2] | The concentration of PDGF-BB:VEGFR2 complex |
| [PDBB:PDRa] | The concentration of PDGF-BB:PDGFRα complex |
| [PDBB:PDRb] | The concentration of PDGF-BB:PDGFRβ complex |
| [R1:N1] | The concentration of VEGFR1:NRP1 complex |

**Chemical reactions**

- **VEGF-A binding to receptors**

$$VA+\overset{kon_{VA,R1}}{R1\overset{\leftrightarrow}{koff_{VA,R1}}} VA:R1$$

$$VA+\overset{kon_{VA,R2}}{R2\overset{\leftrightarrow}{koff_{VA,R2}}} VA:R2$$

$$VA+\overset{kon_{VA,N1}}{N1\overset{\leftrightarrow}{koff_{VA,N1}}} VA:N1$$

$$VA+\overset{kon_{VA,PDRa}}{PDRa\overset{\leftrightarrow}{koff_{VA,PDRa}}} VA:PDRa$$

$$VA+\overset{kon_{VA,PDRb}}{PDRb\overset{\leftrightarrow}{koff_{VA,PDRb}}} VA:PDRb$$

- **VEGF-B binding to receptors**

$$VB+\overset{kon_{VB,R1}}{R1\overset{\leftrightarrow}{koff_{VB,R1}}} VB:R1$$

$$VB+\overset{kon_{VB,R1}}{N1\overset{\leftrightarrow}{koff_{VB,R1}}} VB:N1$$

- **PlGF binding to receptors**

$$Pl+\overset{kon_{VB,R1}}{R1\overset{\leftrightarrow}{koff_{VB,R1}}} Pl:R1$$

$$Pl+\overset{kon_{Pl,R1}}{N1\overset{\leftrightarrow}{koff_{Pl,R1}}} Pl:N1$$

- **PDGF-AA binding to receptors**

$$PDAA+\overset{kon_{PDAA,R2}}{R2\overset{\leftrightarrow}{koff_{PDAA,R2}}} PDAA:R2$$

$$PDAA+\overset{kon_{PDAA,PDRa}}{PDRa\overset{\leftrightarrow}{koff_{PDAA,PDRa}}} PDAA:PDRa$$

- **PDGF-AB binding to receptors**

$$PDAB+\overset{kon_{PDAB,R2}}{R2\overset{\leftrightarrow}{koff_{PDAB,R2}}} PDAB:R2$$

$$PDAB+\overset{kon_{PDAB,PDRa}}{PDRa\overset{\leftrightarrow}{koff_{PDAB,PDRa}}} PDAB:PDRa$$

$$PDAB+\overset{kon_{PDAB,PDRb}}{PDRb\overset{\leftrightarrow}{koff_{PDAB,PDRb}}} PDAB:PDRb$$

- **PDGF-BB binding to receptors**

$$PDBB+\overset{kon_{PDBB,R2}}{R2\overset{\leftrightarrow}{koff_{PDBB,R2}}} PDBB:R2$$

$$PDBB+\overset{kon_{PDBB,PDRa}}{PDRa\overset{\leftrightarrow}{koff_{PDBB,PDRa}}} PDBB:PDRa$$

$$PDBB+\overset{kon_{PDBB,PDRb}}{PDRb\overset{\leftrightarrow}{koff_{PDBB,PDRb}}} PDBB:PDRb$$

- **Receptor coupling**

$$VA:R2+\overset{kc_{VA:R2,N1}}{N1\overset{\leftrightarrow}{koff_{VA:R2,N1}}} VA:R2:N1$$

$$VA:N1+\overset{kc_{VA:N1,R2}}{R2\overset{\leftrightarrow}{koff_{VA:N1,R2}}} VA:R2:N1$$

$$R1+\overset{kon_{R1,N1}}{N1\overset{\leftrightarrow}{koff_{R1,N1}}} R1:N1$$

**Ordinary differential equations**

$$\frac{d[VA]}{dt}=-kon_{VA,R1}\left[ VA \right]\left[ R1 \right]+koff_{VA,R1}\left[ VA:R1 \right]-kon_{VA,R2}\left[ VA \right]\left[ R2 \right]+koff_{VA,R2}\left[ VA:R2 \right]-kon_{VA,N1}\left[ VA \right]\left[ N1 \right]+koff_{VA,N1}\left[ VA:N1 \right]-kon_{VA,PDRa}\left[ VA \right]\left[ PDRa \right]+koff_{VA,PDRa}\left[ VA:PDRa \right]-kon_{VA,PDRb}\left[ VA \right]\left[ PDRb \right]+koff_{VA,PDRb}[VA:PDRb]$$

$$\frac{d\left[ VB \right]}{dt}=-kon_{VB,R1}\left[ VB \right]\left[ R1 \right]+koff_{VB,R1}\left[ VB:R1 \right]-kon_{VB,N1}\left[ VB \right]\left[ N1 \right]+koff_{VB,N1}[VB:N1]$$

$$\frac{d\left[ Pl \right]}{dt}=-kon_{Pl,R1}\left[ Pl \right]\left[ R1 \right]+koff_{Pl,R1}\left[ Pl:R1 \right]-kon_{Pl,N1}\left[ Pl \right]\left[ N1 \right]+koff_{Pl,N1}[Pl:N1]$$

$$\frac{d\left[ PDAA \right]}{dt}=-kon_{PDAA,R2}\left[ PDAA \right]\left[ R2 \right]+koff_{PDAA,R2}\left[ PDAA:R2 \right]-kon_{PDAA,PDRa}\left[ PDAA \right]\left[ PDRa \right]+koff_{PDAA,PDRa}\left[ PDAA:PDRa \right]$$

$$\frac{d\left[ PDAB \right]}{dt}=-kon_{PDAB,R2}\left[ PDAB \right]\left[ R2 \right]+koff_{PDAB,R2}\left[ PDAB:R2 \right]-kon_{PDAB,PDRa}\left[ PDAB \right]\left[ PDRa \right]+koff_{PDAB,PDRa}\left[ PDAB:PDRa \right]-kon_{PDAB,PDRb}\left[ PDAB \right]\left[ PDRb \right]+koff_{PDAB,PDRb}\left[ PDAB:PDRb \right]$$

$$\frac{d\left[ PDBB \right]}{dt}=-kon_{PDBB,R2}\left[ PDBB \right]\left[ R2 \right]+koff_{PDBB,R2}\left[ PDBB:R2 \right]-kon_{PDBB,PDRa}\left[ PDBB \right]\left[ PDRa \right]+koff_{PDBB,PDRa}\left[ PDBB:PDRa \right]-kon_{PDBB,PDRb}\left[ PDBB \right]\left[ PDRb \right]+koff_{PDBB,PDRb}[PDBB:PDRb]$$

$$\frac{d\left[ R1 \right]}{dt}=-kon_{VA,R1}\left[ VA \right]\left[ R1 \right]+koff_{VA,R1}\left[ VA:R1 \right]-kon_{VB,R1}\left[ VB \right]\left[ R1 \right]+koff_{VB,R1}\left[ VB:R1 \right]-kon_{Pl,R1}\left[ Pl \right]\left[ R1 \right]+koff_{Pl,R1}\left[ Pl:R1 \right]-kon_{R1,N1}\left[ R1 \right]\left[ N1 \right]+koff_{R1,N1}\left[ R1:N1 \right]$$

$$\frac{d\left[ R2 \right]}{dt}=-kon_{VA,R2}\left[ VA \right]\left[ R2 \right]+koff_{VA,R2}\left[ VA:R2 \right]-kc_{VAN1,R2}\left[ VA:N1 \right]\left[ R2 \right]+koff_{VAN1,R2}\left[ VA:R2:N1 \right]-kon_{PDAA,R2}\left[ PDAA \right]\left[ R2 \right]+koff_{PDAA,R2}\left[ PDAA:R2 \right]-kon_{PDAB,R2}\left[ PDAB \right]\left[ R2 \right]+koff_{PDAB,R2}\left[ PDAB:R2 \right]-kon_{PDBB,R2}\left[ PDBB \right]\left[ R2 \right]+koff_{PDBB,R2}\left[ PDBB:R2 \right]$$

$$\frac{d\left[ N1 \right]}{dt}=-kon_{VA,N1}\left[ VA \right]\left[ N1 \right]+koff_{VA,N1}\left[ VA:N1 \right]-kc_{VAR2,N1}\left[ VA:R2 \right]\left[ N1 \right]+koff_{VAR2,N1}[VA:R2:N1]-kon_{VB,N1}\left[ VB \right]\left[ N1 \right]+koff_{VB,N1}[VB:N1]-kon_{Pl,N1}\left[ Pl \right]\left[ N1 \right]+koff_{Pl,N1}[Pl:N1]-kon_{R1,N1}\left[ R1 \right]\left[ N1 \right]+koff_{R1,N1}\left[ R1:N1 \right]$$

$$\frac{d\left[ PDRa \right]}{dt}=-kon_{VA,PDRa}\left[ VA \right]\left[ PDRa \right]+koff_{VA,PDRa}\left[ VA:PDRa \right]-kon_{PDAA,PDRa}\left[ PDAA \right]\left[ PDRa \right]+koff_{PDAA,PDRa}\left[ PDAA:PDRa \right]-kon_{PDAB,PDRa}\left[ PDAB \right]\left[ PDRa \right]+koff_{PDAB,PDRa}\left[ PDAB:PDRa \right]-kon_{PDBB,PDRa}\left[ PDBB \right]\left[ PDRa \right]+koff_{PDBB,PDRa}\left[ PDBB:PDRa \right]$$

$$\frac{d\left[ PDRb \right]}{dt}=-kon_{VA,PDRb}\left[ VA \right]\left[ PDRb \right]+koff_{VA,PDRb}[VA:PDRb]-kon_{PDAB,PDRb}\left[ PDAB \right]\left[ PDRb \right]+koff_{PDAB,PDRb}\left[ PDAB:PDRb \right]-kon_{PDBB,PDRb}\left[ PDBB \right]\left[ PDRb \right]+koff_{PDBB,PDRb}[PDBB:PDRb]$$

$$\frac{d\left[ VA:R1 \right]}{dt}=kon_{VA,R1}\left[ VA \right]\left[ R1 \right]-koff_{VA,R1}\left[ VA:R1 \right]$$

$$\frac{d\left[ VA:R2 \right]}{dt}= kon_{VA,R2}\left[ VA \right]\left[ R2 \right]-koff_{VA,R2}\left[ VA:R2 \right]+kc_{VAR2,N1}\left[ VA:R2 \right]\left[ N1 \right]-koff_{VAR2,N1}[VA:R2:N1]$$

$$\frac{d\left[ VA:N1 \right]}{dt}=kon_{VA,N1}\left[ VA \right]\left[ N1 \right]-koff_{VA,N1}\left[ VA:N1 \right]+kc_{VAN1,R2}\left[ VA:N1 \right]\left[ R2 \right]-koff_{VAN1,R2}\left[ VA:R2:N1 \right]$$

$$\frac{d\left[ VA:R2:N1 \right]}{dt}=kc_{VAN1,R2}\left[ VA:N1 \right]\left[ R2 \right]-koff_{VAN1,R2}\left[ VA:R2:N1 \right]+kc_{VAR2,N1}\left[ VA:R2 \right]\left[ N1 \right]-koff_{VAR2,N1}[VA:R2:N1]$$

$$\frac{d\left[ VA:PDRa \right]}{dt}=kon_{VA,PDRa}\left[ VA \right]\left[ PDRa \right]-koff_{VA,PDRa}\left[ VA:PDRa \right]$$

$$\frac{d\left[ VA:PDRb \right]}{dt}=kon_{VA,PDRb}\left[ VA \right]\left[ PDRb \right]-koff_{VA,PDRb}[VA:PDRb]$$

$$\frac{d\left[ VB:R1 \right]}{dt}=kon_{VB,R1}\left[ VB \right]\left[ R1 \right]-koff_{VB,R1}\left[ VB:R1 \right]$$

$$\frac{d\left[ VB:N1 \right]}{dt}= kon_{VB,N1}\left[ VB \right]\left[ N1 \right]-koff_{VB,N1}[VB:N1]$$

$$\frac{d\left[ Pl:R1 \right]}{dt}=kon_{Pl,R1}\left[ Pl \right]\left[ R1 \right]-koff_{Pl,R1}\left[ Pl:R1 \right]$$

$$\frac{d\left[ Pl:N1 \right]}{dt}= kon_{Pl,N1}\left[ Pl \right]\left[ N1 \right]-koff_{Pl,N1}[Pl:N1]$$

$$\frac{d\left[ PDAA:R2 \right]}{dt}=kon_{PDAA,R2}\left[ PDAA \right]\left[ R2 \right]+koff_{PDAA,R2}\left[ PDAA:R2 \right]$$

$$\frac{d\left[ PDAA:PDRa \right]}{dt}=kon_{PDAA,PDRa}\left[ PDAA \right]\left[ PDRa \right]-koff_{PDAA,PDRa}\left[ PDAA:PDRa \right]$$

$$\frac{d\left[ PDAB:R2 \right]}{dt}=kon_{PDAB,R2}\left[ PDAB \right]\left[ R2 \right]-koff_{PDAB,R2}\left[ PDAB:R2 \right]$$

$$\frac{d\left[ PDAB:PDRa \right]}{dt}= kon_{PDAB,PDRa}\left[ PDAB \right]\left[ PDRa \right]-koff_{PDAB,PDRa}\left[ PDAB:PDRa \right]$$

$$\frac{d\left[ PDAB:PDRb \right]}{dt}=kon_{PDAB,PDRb}\left[ PDAB \right]\left[ PDRb \right]-koff_{PDAB,PDRb}\left[ PDAB:PDRb \right]$$

$$\frac{d\left[ PDBB:R2 \right]}{dt}=kon_{PDBB,R2}\left[ PDBB \right]\left[ R2 \right]-koff_{PDBB,R2}\left[ PDBB:R2 \right]$$

$$\frac{d\left[ PDBB:PDRa \right]}{dt}=kon_{PDBB,PDRa}\left[ PDBB \right]\left[ PDRa \right]-koff_{PDBB,PDRa}\left[ PDBB:PDRa \right]$$

$$\frac{d\left[ PDBB:PDRb \right]}{dt}=kon_{PDBB,PDRb}\left[ PDBB \right]\left[ PDRb \right]-koff_{PDBB,PDRb}[PDBB:PDRb]$$

$$\frac{d\left[ R1:N1 \right]}{dt}=kon_{R1,N1}\left[ R1 \right]\left[ N1 \right]-koff_{R1,N1}\left[ R1:N1 \right]$$
